# Mono- and bi-allelic protein truncating variants in alpha-actinin 2 cause cardiomyopathy through distinct mechanisms

**DOI:** 10.1101/2020.11.19.389064

**Authors:** Malene E Lindholm, David Jimenez-Morales, Han Zhu, Kinya Seo, David Amar, Chunli Zhao, Archana Raja, Roshni Madhvani, Cedric Espenel, Shirley Sutton, Colleen Caleshu, Gerald J Berry, Kara S. Motonaga, Kyla Dunn, Julia Platt, Euan A Ashley, Matthew T Wheeler

## Abstract

Alpha-actinin 2 (ACTN2) anchors actin within cardiac sarcomeres. The mechanisms linking *ACTN2* mutations to myocardial disease phenotypes are unknown. Here, we characterize patients with novel *ACTN2* mutations to reveal insights into the physiological function of ACTN2. Patient-derived iPSC-cardiomyocytes harboring ACTN2 protein-truncating variants were hypertrophic, displayed sarcomeric structural disarray, impaired contractility, and aberrant Ca^2+^-signaling. In heterozygous indel cells, the truncated protein incorporates into cardiac sarcomeres, leading to aberrant Z-disc structure. In homozygous stop-gain cells, affinity-purification mass spectrometry reveals an intricate ACTN2 interactome with sarcomere and sarcolemma-associated proteins. Loss of the C-terminus of ACTN2 disrupts interaction with ACTN1 and GJA1, two sarcolemma-associated proteins, that may lead to the clinical arrhythmic and relaxation defects. The causality of the stop-gain mutation was verified using CRISPR-Cas9 gene editing. Together, these data advance our understanding of the role of ACTN2 in the human heart and establish recessive inheritance of *ACTN2* truncation as causative of disease.

## Introduction

Inherited cardiomyopathy and arrhythmia syndromes are common indications for heart transplantation and leading causes of sudden cardiac death in children and adolescents. Clinical genetic testing of families with atypical presentations often leaves clinicians and families with variants of uncertain significance. Growing evidence has shown a diverse genetic etiology of early-onset cardiomyopathies including dilated, hypertrophic, restrictive, and arrhythmogenic cardiomyopathies due to mutations of genes of the sarcomere and the sarcolemma^1–3^.

Alpha-actinin 2 *(ACTN2)* is an integral sarcomeric protein known to cross-link sarcomeric actin and titin filaments in the Z-disc of cardiac- and skeletal-myocyte sarcomeres. It is primarily expressed in cardiac and skeletal muscle^4, 5^, where it forms an antiparallel homodimer, with an N-terminal actin-binding domain, a central rod region with spectrin-like repeats, and a C-terminal titin-binding region containing a calmodulin-like and EF-hand (helix-loop-helix) domain^6^. Prior mechanistic work has shown multiple functions for alpha-actinin 2, including anchoring of proteins within the sarcomere^7–10^, regulation of ion channels^11,12^, and indirect control of striated muscle gene expression through enhanced transactivation activity of nuclear receptors^13^. A missense mutation in *ACTN2* was first identified in a patient with dilated cardiomyopathy (DCM) in 2003^14^. Since then, additional studies have reported heterozygous missense mutations in *ACTN2* that co-segregate with both dilated and hypertrophic cardiomyopathy (HCM)^15–21^. Common genetic variation in a regulatory element modulating *ACTN2* expression was recently associated with heart failure with reduced ejection fraction^22^. Despite these association studies, there is limited evidence for *ACTN2* as a causative gene for cardiomyopathy^23, 24^. In addition, very few studies have found truncating variants in *ACTN2*, without evidence of clear genetic association^*25-27*^, and the relationship of truncating variants with cardiac disease manifestations has not been established.

To date, only autosomal dominant inheritance of *ACTN2* variants has been described in association with striated muscle disease and there is very limited mechanistic data on how *ACTN2* mutations cause disease. Herein, we demonstrate how structurally aberrant ACTN2 causes cardiac disease through two distinct mechanisms; one through integration of the truncated protein into the cardiac sarcomere, and the second through loss of integral protein-protein interactions. We describe these genetic variations, resulting in truncated ACTN2 proteins^28^, as causative of cardiomyopathy with prominent arrhythmic phenotypes in the heterozygous state, and as causative of a progressive, severe restrictive cardiomyopathy when in the homozygous state. Moreover, our successful modeling of *ACTN2* genetic disease using hiPSC-CMs presents an opportunity to identify targeted therapies, which are especially pertinent for this young patient population.

## Results

To explore if structural variation in ACTN2 is causative of early-onset cardiomyopathy, we identified patients in the Stanford Center for Inherited Cardiovascular Disease database with rare or novel *ACTN2* variants using a custom mutation pipeline optimized for rare variant discovery. The first patient showed homozygosity for a novel variant in *ACTN2*, p.Gln860Stop (Q860X), which is absent in gnomAD^29^. The patient presented in infancy with symptoms of tachypnea, reduced ejection fraction, as well as low muscle bulk and tone (pedigree presented in **Figure 1A**). Due to conflicting data regarding the etiology of her cardiomyopathy, she underwent diagnostic endomyocardial biopsy at age 16, which showed non-specific patchy hypertrophy and fine interstitial fibrosis, but no myofibrillar loss nor Z-band remnants (**Figure 1G**, **Supplementary Figure 1)**. Early in the third decade of life, the patient developed progressive heart failure symptoms and episodes of atrial fibrillation with rapid ventricular response. Echocardiogram at that time was notable for severe biatrial enlargement, moderate biventricular dysfunction, and diastolic dysfunction (**Figure 1F**); right heart catheterization showed restrictive cardiomyopathy (RCM) with equalization of ventricular filling pressures. She underwent orthotopic heart transplantation at age 23. Explant ventricular myocardial tissue sections showed mild hypertrophy and severe interstitial fibrosis (**Figure 1G**). To verify the sequence-identified *ACTN2* truncation and investigate possible nonsense-mediated decay, we isolated protein from the explanted left ventricle, right ventricle and from a contemporaneous biopsy of intercostal skeletal muscle. Western blot analysis showed that protein levels of ACTN2 were normal in patient cardiac tissue compared to corresponding cardiac tissue from healthy control individuals, and confirmed the sequence-based predicted size reduction of approximately 4kDa (**Figure 1H**). We further characterized the patient’s left ventricular tissue through RNA sequencing (**Supplementary Figure 2**, see **Materials and Methods** for details). Gene set enrichment analysis showed positive enrichment of extracellular matrix remodeling, collagen biosynthesis, and mRNA splicing gene sets (**Supplementary Figure 2**). Highly ranked genes included collagens (*e.g. COL1A1, COL1A2*, and *COL4A1)*, fibronectin *(FN1)* and transforming growth factor β (*TGFB1*), that have been shown to be involved in pathologic cardiomyocyte hypertrophy and fibrosis.^30^ These findings were highly indicative of the fibrosis observed in the patient cardiac tissue. Common markers of pathologic cardiomyocyte hypertrophy and cardiomyopathy, *e.g. NPPA* and *NPPB*, were also elevated. Although ACTN2 protein levels were similar in the patient compared to controls, ACTN2 transcription was higher in the patient (**Figure 1I**). To evaluate if ACTN2 is differentially expressed in DCM and heart failure, we compared ACTN2 expression levels in the left ventricle of 25 healthy individuals, 24 DCM patients, and 21 patients with ischemic heart disease from the MAGNet consortium^31, 32^. No difference in ACTN2 expression was observed in these samples (**Supplementary Figure 3**).

**Figure 1.**
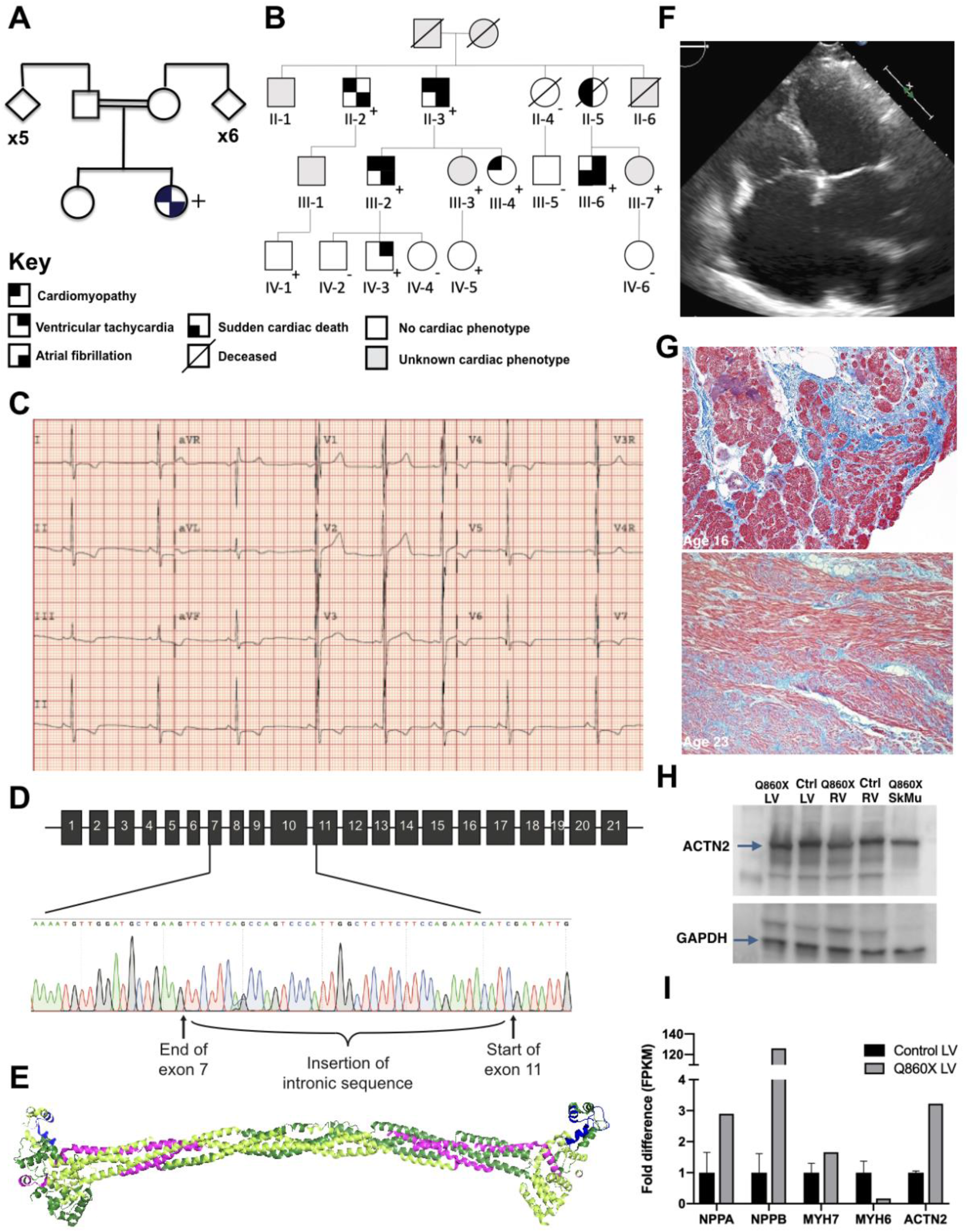
Protein-truncating alpha-actinin 2 variants cause diverse cardiac symptoms. A) Pedigree for a patient homozygous for a C-terminal truncation of ACTN2 (Q860X). B) Pedigree for a family heterozygous for a large indel in ACTN2 (Del). C) ECG recordings of the Del family proband shows T-wave inversion. D) Sanger sequencing showed insertion of a 41bp intronic sequence in addition to the exon 8-10 deletion in the Del family, producing an inframe indel. E) The protein-level truncating variants displayed in the 3D structure of ACTN2. The Q860X missing C-terminal region is shown in blue, while the exon 8-10 deletion of the indel is shown in pink. F) Transthoracic echocardiography of Q860X showed restrictive cardiomyopathy. G) Trichrome staining of an endomyocardial biopsy obtained at age 16 (top) and the explanted left ventricle tissue from transplant at age 23 showed coarse and fine interstitial fibrosis of cardiomyopathy. H) Western blot analysis of ACTN2 protein expression in the cardiac tissue indicated no nonsense-mediated decay, and confirmed the predicted size reduction of the protein by 4kDa. I) RNA sequencing analysis of the Q860X patient’s left ventricle compared to healthy human control hearts (n=3) showed elevated expression of known markers of hypertrophy *(NPPA* and *NPPB)*, as well as an elevated expression of *ACTN2* itself.

The second identified proband (Del 1) presented at age 16 with an episode of exertional syncope (individual IV-3 in **Figure 1B**). ECG showed T-wave inversion and repolarization abnormalities (**Figure 1C**). Clinical panel genetic testing identified a heterozygous novel 4.3kb deletion in *ACTN2* of exons 8 through 10, (chr 1: 236,898,807-236,903,093) classified as a variant of unknown significance. A comprehensive 5-generation family history revealed diverse cardiac symptoms, including two early sudden cardiac deaths, ventricular tachyarrhythmias, atrial fibrillation, left ventricular noncompaction, diastolic dysfunction and symptomatic heart failure. Subsequent clinical screening and single locus testing of 1^st^ degree relatives of affected individuals identified six additional family members with manifestations of ventricular conduction delay on ECG, early onset atrial arrhythmia, left ventricular noncompaction cardiomyopathy, and/or symptoms consistent with early onset heart failure with preserved ejection fraction, each of whom harbored the *ACTN2* variant. Samples were also contributed by the proband’s father, who exhibited extensive T-wave inversions on ECG, atrial fibrillation, and nonsustained ventricular tachycardia (Del 2). The deletion identified through clinical genetic testing in Family 2 was verified using Sanger sequencing, which identified a 41bp intronic insertion from a central region of intron 10 of ACTN2, in addition to the 4.3 kb deletion (**Figure 1D**). The resulting sequence is an in-frame deletion that would translate into an internally truncated protein of 771 residues compared to the full-length ACTN2 of 894 amino acids. The deleted region is shown in pink in the structural model of ACTN2 (**Figure 1E**).

Genetic variants in *ACTN2* are rare and the gene is intolerant to loss of function variation (pLI score of 1.0)^29^. The number of unique exonic variants in GnomAD was 506 after removal of synonymous variants (search: Nov 12 2020). The distribution of variants per exon is illustrated in **Supplementary Figure 4**. ClinVar^33^ reports 48 pathogenic variants (search: Nov 12 2020); 42 copy number variants, 4 missense, 1 splice acceptor and 1 nonsense variant, which belongs to the first patient included in this study.

### Patient-derived hiPSC-cardiomyocytes display elevated expression of metabolic genes

To functionally model the disease caused by truncated ACTN2 proteins, we utilized patient-derived and healthy control human induced pluripotent stem cells (hiPSCs) differentiated into cardiomyocytes (hiPSC-CMs). Pluripotency of the hiPSC lines was confirmed through immunofluorescence staining for the pluripotency-markers Nanog, Oct4, SSEA3 and TRA-1-81 (**Supplementary Figure 5**). ACTN2 expression occurred at day 6 post initiation of differentiation (**Supplementary Figure 5**). All subsequent experiments were performed ~30 days post initiation of differentiation. We started by performing RNA sequencing on the three patient hiPSC-CMs and two different control hiPSC-CMs. Basic features of the analysis are shown in **Supplementary Figure 6**. Expression of *ACTN2* was somewhat higher in both Q860X (logFC 0.8) and the heterozygous patients (logFC 1.14). Interestingly, in all patients, the most elevated gene was *MYL2* (Q860X: logFC 5.23, Del: logFC 4.90), coding for Myosin light chain 2, a key contractile cardiac protein. We performed a gene set enrichment analysis to investigate the general pathways affected in disease. Multiple pathways were common between the two genetic conditions; a Venn diagram summarizing this is shown in **Supplementary Figure 6E**. Notably, induction of metabolic pathways was a common feature, with respiratory electron transport and gluconeogenesis represented among the top five enriched pathways. Cristae formation was another common pathway, that is mainly driven by ATP synthase related genes. Among the top down-regulated pathways, only keratinization was common between the two genetic conditions. This is mainly driven by kallikreins, which are serine phosphatases with diverse functions. In Q860X, receptor-type tyrosine-protein phosphatases was the top down-regulated pathway. This was driven by lower expression of several SLIT and NTRK Like Family Members *(SLITRK* genes); integral transmembrane proteins expressed in the heart but with unknown cardiac function. In the patients with the heterozygous indel, lower expression level was seen for many structural genes associated with extracellular matrix organization and collagen formation. Multiple different collagens *(e.g COL1A1, COL3A1, COL9A3* and *COL6A3)* and genes associated with collagen assembly *(DCN, LUM* and *LOX)* are represented.

To verify that the pathology is not due to lower ACTN2 protein levels, protein expression was investigated using Western Blot. There was no evidence of proteolytic or ER mediated degradation of the aberrant protein due to Q860X; protein levels were similar to controls. Interestingly, we found that the exon 8-10 indel allele also produces a protein product (**Figure 2A-B**), although the levels were lower compared to the full length protein in the same samples. RNA sequencing data from Del 1 and Del 2 similarly showed evidence of limited nonsense mediated decay of the truncated transcript with relative expression of 20% and 30% respectively, in relation to the full length transcript (**Supplementary Figure 7**).

**Figure 2.**
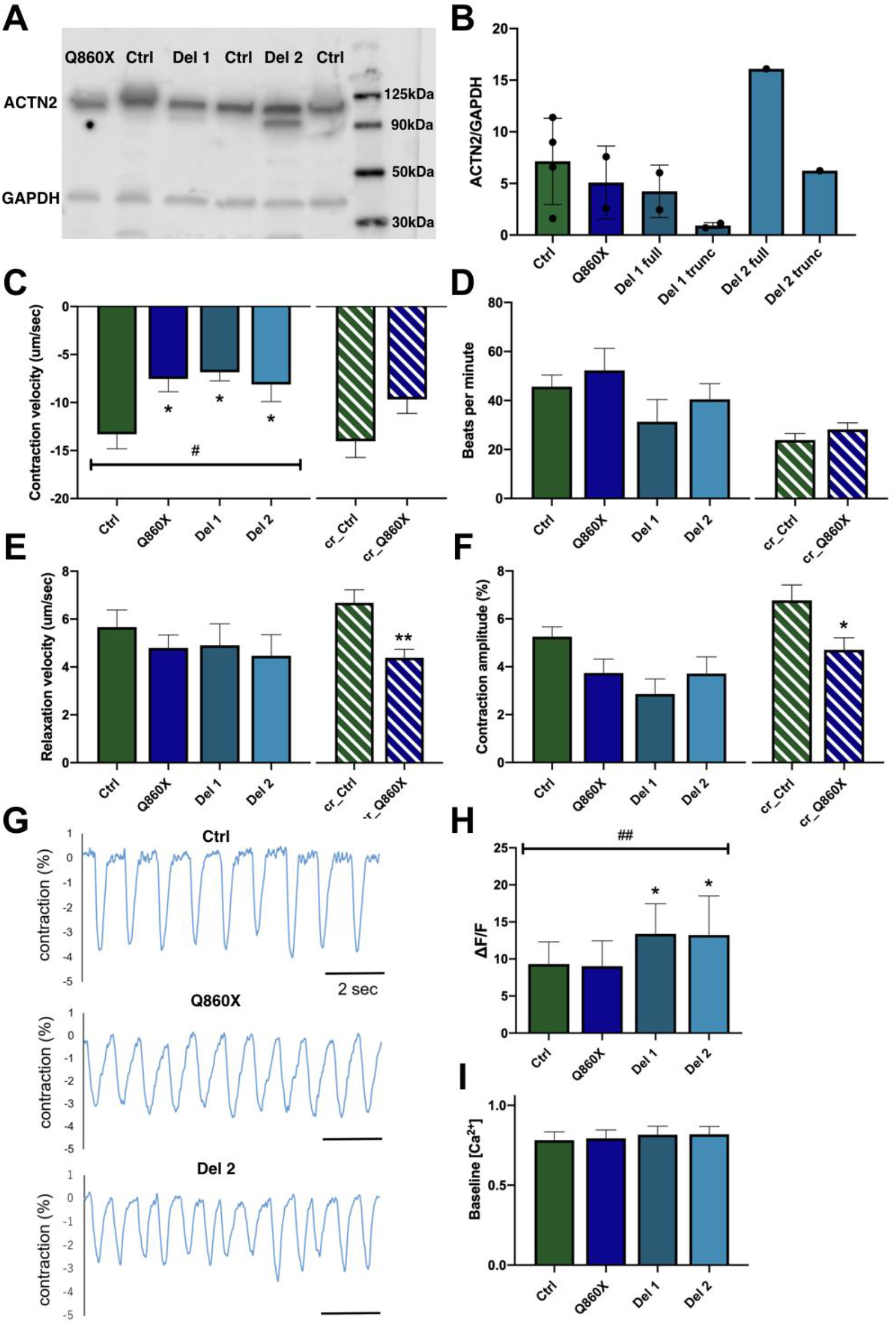
ACTN2 truncation causes contractile dysfunction. A-B) Western blot analysis showed that hiPSC-CMs from Q860X, Del 1 and Del 2 expressed ACTN2. Del 1 and Del 2 also expressed the truncated protein product, although at a lower level compared to the full length protein. C-F) Contractility analysis of single-cell hiPSC-CMs using video-based edge detection through the IonOptix system. G) Representative traces from the contractile analysis. H-I) Ca^2+^-handling analysis of single cell hiPSC-CMs. Cells were loaded with a ratiometric Ca^2+^ probe and analyzed using video-based edge detection through the IonOptix system. * indicates significant difference compared to the control group at *P*<0.05 using a one-way ANOVA with Dunnett’s multiple comparisons test.

### Truncated ACTN2 causes contractile dysfunction

Next, we sought to assess the functional implications of a truncated ACTN2 protein. We utilized video-based edge detection to measure the contractile function of single hiPSC-CMs. A significantly lower contractile velocity was found in all three patient lines, with no difference in relaxation velocity (**Figure 2C-D**). There was no difference in beating frequency (**Figure 2E**) or contraction amplitude (**Figure 2F**), which measures the degree of cellular shortening during each contraction. Representative traces from the contractility analysis is shown in **Figure 2G**. To confirm that the C-terminal truncation (Q860X) is the causal disease variant, we used CRISPR-Cas9 to introduce the truncation into healthy hiPSCs to investigate if that would recapitulate the disease phenotype. Two different guide RNAs were introduced together with the Cas9 to cut out the C-terminal region of both copies of *ACTN2.* Sequencing and gel-based size verification were used to confirm successful gene editing (see **Materials and Methods** and **Supplementary Figure 8** for details). After differentiation into hiPSC-CMs, single-cell contractility analysis was performed for the CRISPR:ed control line that mirrored the disease line (cr_Q860X) compared to the isogenic healthy control line (cr_Ctrl). Although there was no significant reduction in contractile velocity, there was a similar trend (p=0.058) to what was observed for Q860X (**Figure 2C**). The differences in relaxation velocity (**Figure 2D**) and contraction amplitude (**Figure 2F**) for cr_Q860X was also very similar to Q860X, while the changes were statistically significant for cr_Q860X.

Next, because cellular Ca^2+^ handling is key for cardiomyocyte contractile function, we investigated if this was altered as a consequence of structurally aberrant ACTN2. We loaded live hiPSC-CMs with a ratiometric Ca^2+^-binding fluorescent probe and Ca^2+^ signals were evaluated using the IonOptix system. Interestingly, both patients with the heterozygous indel (Del 1 and Del 2) showed higher intracellular Ca^2+^ signalling (ΔF/F) compared to controls, while there was no effect in the patient with the homozygous truncation (Q860X) (**Figure 2H**). This suggests that the contractile dysfunction is at the sarcomeric level for the heterozygous family, where elevated Ca^2+^ signalling fails to compensate for the low contractility. There was no difference in baseline Ca^2+^ levels (**Figure 2I**).

### ACTN2 dysfunction induces cardiomyocyte hypertrophy

In response to the hypertrophic phenotype of the patients and the altered cardiac contractility observed, which commonly leads to hypertrophy, we continued by analyzing cell size in the hiPSC-CMs. Single hiPSC-CMs were fixed and fluorescently co-labelled with ACTN2 as a cardiomyocytespecific marker and an actin probe. After imaging, cell edges were traced and analyzed blinded to disease status. Representative images for each patient line and a healthy control line are shown in **Figure 3A**. Patient cells were hypertrophic compared to healthy control cells (**Figure 3B**), which was also observed for the CRISPR:ed line (cr_Q860X, **Supplementary Figure 9**). To further determine the impact of a disrupted ACTN2 function on development of a hypertrophic phenotype, we used a knock down approach in neonatal rat ventricular myocytes (NRVMs). The results showed that cells targeted with siRNA against ACTN2 displayed marked hypertrophy compared to cells treated with a negative control siRNA (**Supplementary Figure 9**).

**Figure 3.**
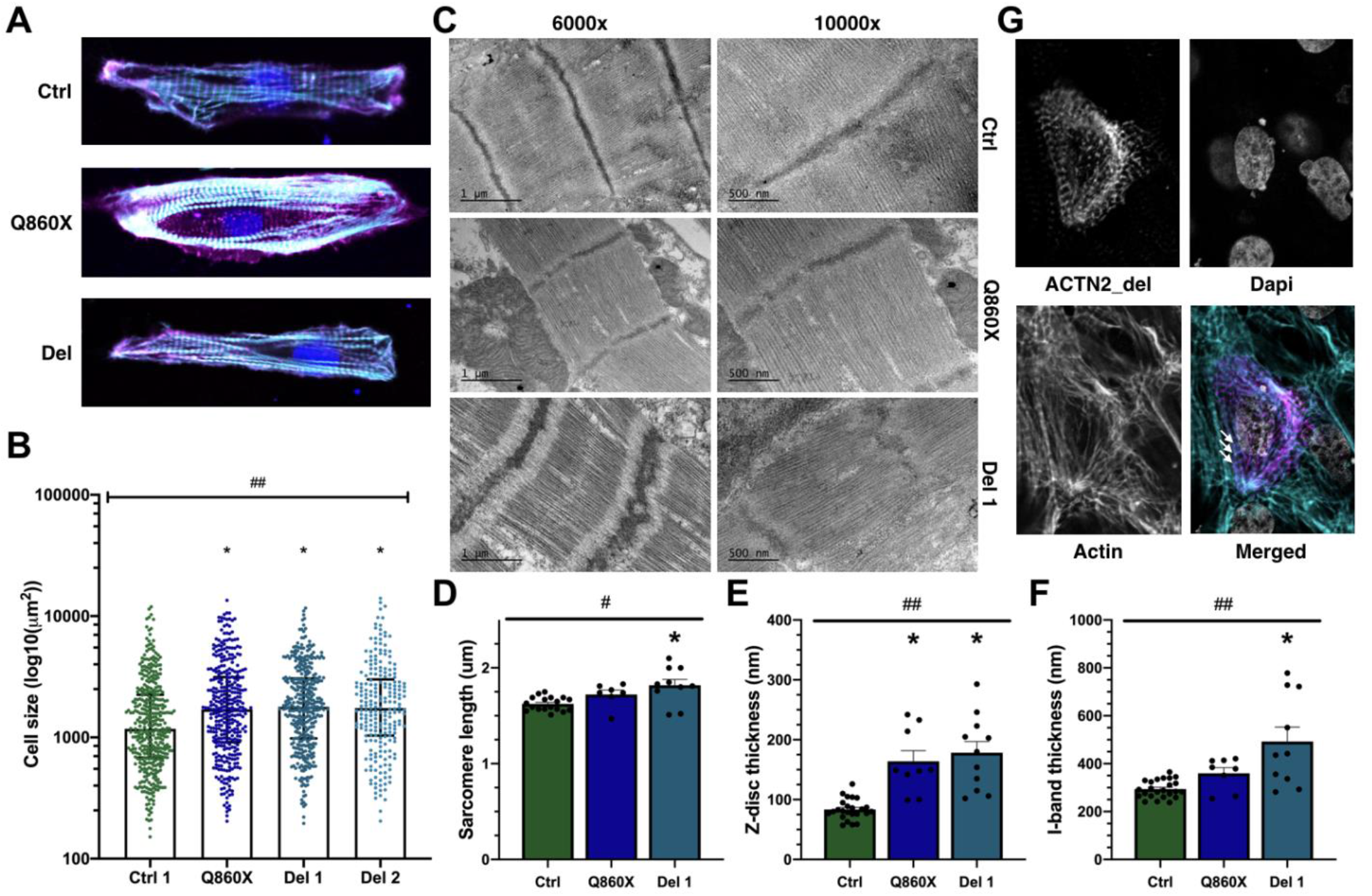
Patient-specific hiPSC-CMs are hypertrophic and display sarcomeric structural disarray. A) Representative hiPSC-CMs from each line, with ACTN2 shown in purple, actin in turquoise and nuclei in blue (Dapi). B) Cell size of single-cell hiPSC-CMs. C) Representative transmission electron microscopy images of control hiPSC-CMs, Q860X and Del 1 at two different magnifications (6000X and 10000X). D-F) Quantitative analysis of the electron microscopy images, showing data for sarcomere length (D), Z-disc thickness (E) and I-band thickness (F). Significant effect from a one-way ANOVA analysis at *P*<0.01 is indicated by #, at *P*<0.0001 by ##. * indicates significant difference compared to control at *P*<0.001 using Dunnett’s multiple comparisons test. G) Representative confocal image of cells transfected with a vector carrying a FLAG-tagged version of ACTN2_del. Image split into the FLAG-tagged ACTN2_del (top left), actin (lower left), Dapi for nuclei (top right), and a merged image with ACTN2_del shown in purple, actin in turquoise and nuclei in grey (lower right). White arrows point to examples where the FLAG-tagged protein carrying the indel has been incorporated into the cardiomyocyte sarcomeres.

### Heterozygous indel in ACTN2 causes sarcomeric structural disarray

The expression of the truncated protein product in the heterozygous family, and the likely sarcomeric cause of the contractile dysfunction lead us to investigate the sarcomeric structure of the cardiomyocytes. Transmission electron microscopy (TEM) of the patient and control hiPSC-CMs revealed severe structural disarray of the sarcomeres (**Figure 3C**). The sarcomeres were somewhat elongated, the Z-discs were larger and the I-bands were thicker compared to healthy control hiPSC-CMs (**Figure 3D-F**). The Z-discs also displayed a zigzag-like shape. Similar structural abnormalities have been observed in skeletal muscle of a patient heterozygous for a missense mutation in exon 18 of *ACTN2*, that was associated with a severe myopathy^34^. The C-terminal truncation in *ACTN2* did not cause any clear structural abnormalities, only an increase in the Z-disc thickness was observed (**Figure 3E**). In zebrafish, a reduction of ACTN2 in cardiac tissue caused increased Z-disc thickness ^35^, supporting that a disrupted *ACTN2* function causes cardiac Z-disc abnormalities to different extent.

The large structural consequence of a heterozygous indel in *ACTN2* in combination with successful translation of the truncated transcript lead us to hypothesize that the pathogenic effect of the indel is due to incorporation of the truncated protein product into the Z-disc. To test this, we transfected a FLAG-tagged ACTN2 transcript, carrying the indel, using a pCMV vector in healthy control hiPSC-CMs (**Supplementary Figure 10**). After fixation and staining with an anti-FLAG antibody, cells were imaged with confocal microscopy. The efficiency of the transfection was low, and some transfected cells were not cardiomyocytes. However, there were multiple successfully transfected hiPSC-CMs, and in accordance with the hypothesis, the protein carrying the large indel was indeed incorporated into the sarcomeres (**Figure 3G, Supplementary Figure 10**). This strongly suggests a dominant negative effect of the *ACTN2* indel, in concordance with the family history.

### C-terminal truncation of ACTN2 disrupts interaction with ACTN1 and GJA1

The normal Ca^2+^ signaling and sarcomere structure in Q860X hiPSC-CMs does not explain the pathogenic contractile dysfunction. Because the patient is homozygous for the C-terminal truncation, we hypothesized that lack of a C-terminal interaction may explain the pathogenic phenotype. To investigate the ACTN2 interactome, we performed co-immunoprecipitation using an ACTN2 monoclonal antibody that had been tested for immunoprecipitation and that was verified to bind to the truncated protein through Western blot. The purified protein complexes were subsequently subjected to LC-MS/MS (Liquid chromatography with tandem mass-spectrometry) (**Figure 4A**). An IgG pulldown was used to control for nonspecific binding.

**Figure 4.**
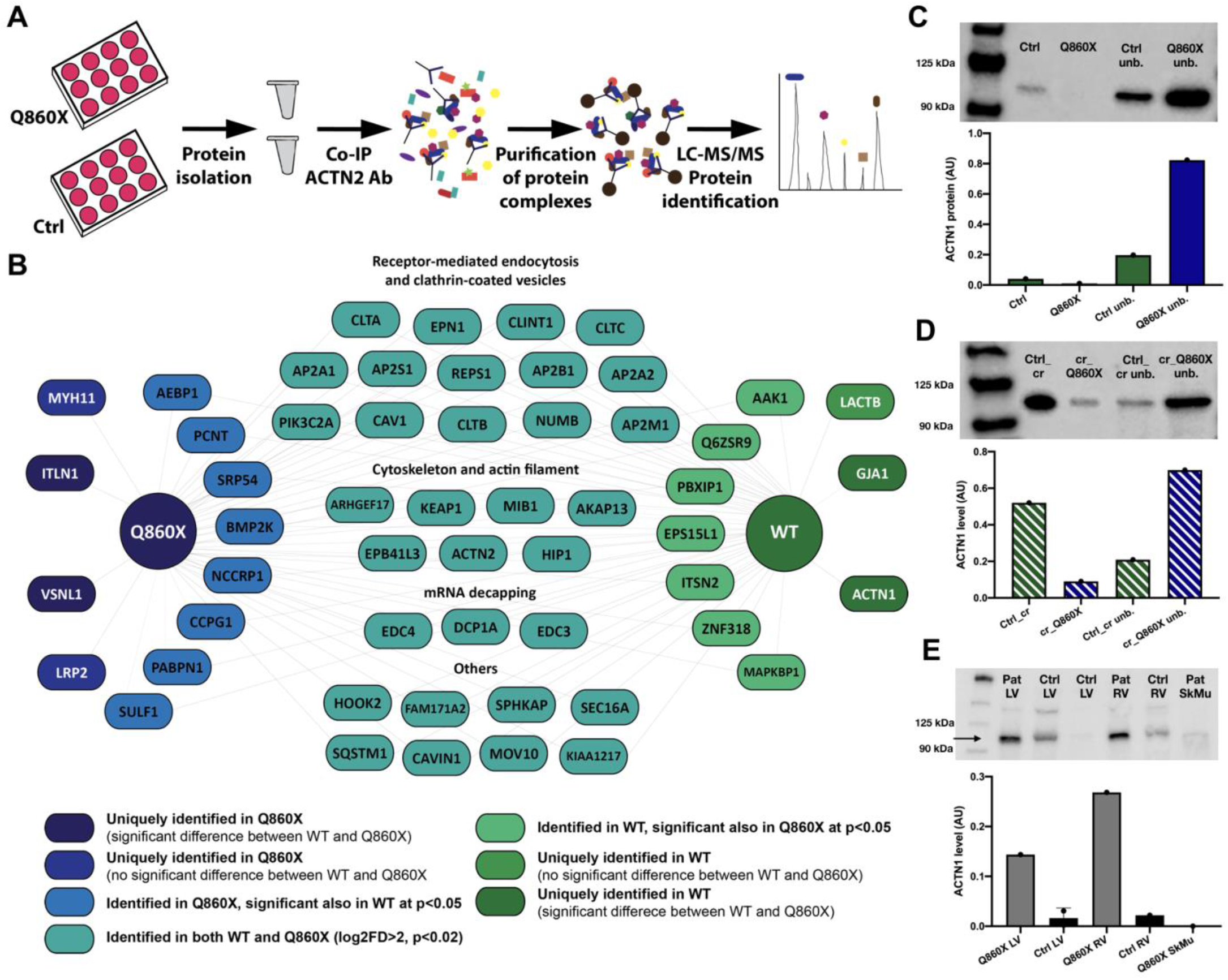
The cardiomyocyte ACTN2 interactome identifies C-terminal region to be necessary for interaction with ACTN1. A) Experimental set-up of the co-immunoprecipitation followed by massspectrometry. IgG pulldown was used as a negative control. B) Analysis of the mass-spectrometry data using MaxQuant identified a common ACTN2 interactome in all hiPSC-CMs in turquoise (iLogFC>2, iPvalue<0.02 in both conditions, at least 3 peptides used for analysis, see **Materials and Methods** for details). Proteins significantly different between conditions and identified only in the WT control (n=3) are shown in dark green, and proteins uniquely identified in Q860X (n=3) are shown in dark blue. C) Co-immunoprecipitation with ACTN2 antibody followed by Western blot using an ACTN1 antibody showed interaction only in healthy control cells compared to patient Q860X, in concordance with the Co-IP mass-spec data. D) Co-immunoprecipitation with ACTN2 antibody followed by Western blot using an ACTN1 antibody showed interaction only in healthy control cells compared to cr_Q860X. E) Expression of ACTN1 in Q860X explanted cardiac left ventricle and right ventricle compared to control.

The interactome of ACTN2 has not been previously investigated in cardiomyocytes or using AP-MS (Affinity-Purification Mass-Spectrometry), while multiple interactors have been identified through cofractionation experiments in cell lines^36^ and through yeast two-hybrid screens^37^, and a few in specific low-throughput affinity-purification experiments in other cell types^38–40^. In human iPSC-CMs we found multiple novel interactors, mainly associated with the cell membrane. Several proteins involved in endocytosis through clathrin-coated pits were significantly pulled down in both healthy WT cells and patient Q860X cells (**Figure 4B**). This demonstrates that ACTN2 plays a role in the cardiac cell membrane, and has a much more intricate regulatory role than what is known to date. Membrane protein extraction with detergent and ultracentrifugation from hiPSC-CMs showed abundant ACTN2 levels in the membrane fraction (**Supplementary Figure 11**). Other studies have demonstrated that ACTN2 is required for the proper membrane localization of a Ca^2+^-activated K^+^-channel^11^, and have shown interactions with membrane-associated proteins, for example Kv1.5 channels ^38^ and Nav1.5 channels^41^. While ACTN2 itself was significantly pulled down as expected, other large structural proteins or their interactors rarely reach significance in pulldown experiments due to the size and abundance of these proteins. Titin (TTN), Actin (ACTC1), Palladin (PALLD), Synaptopodin (SYNPO2) and Myozenin (MYOZ2) are all high abundance sarcomeric proteins that are known interactors of ACTN2^10^, but did not reach significance based on this analysis. Intensity values from mass-spectrometry are not quantitative, but the relative differences between the ACTN2 pulldown and the IgG control indicated higher levels with ACTN2 pulldown compared to the negative IgG control (**Supplementary Figure 11**).

Two proteins that were pulled down with ACTN2 in healthy control hiPSC-CMs, GJA1 and ACTN1, were not identified in the pulldown of the C-terminal truncated ACTN2. GJA1, or Connexin 43, is the main connexin protein in cardiac muscle. It is a gap junction protein important for cardiac electrical conduction^42^. The subcellular location of Connexin 43 in the sarcolemma again suggests an important role for ACTN2 in the membrane. There is no previous evidence of an interaction between ACTN2 and GJA1. ACTN1 is a non-muscle alpha-actinin that is ubiquitously expressed. It has been shown to heterodimerize with the other non-muscle alpha-actinin ACTN4^43^. There are no previous affinity-purification experiments that have shown interaction between ACTN1 and ACTN2, and previous interaction data are from high throughput co-fractionation experiments^36^ and *in vitro* yeast two-hybrid screens^37^. RNA sequencing data showed similar expression levels of the majority of genes identified in the interactome, including ACTN1 and GJA1, showing that this finding is not an effect of differences in expression. Only two identified proteins in the network showed mRNA expression differences, *NCCRP1* and *ITLN1*, both of which were lower in Q860X (**Supplementary Figure 11**).

To corroborate these findings, we repeated the co-immunoprecipitation followed by Western Blot for ACTN1 and GJA1. ACTN1 was identified only in the healthy Ctrl hiPSC-CMs with no visible pulldown in Q860X (**Figure 4C**), in concordance with the mass-spectrometry data. We repeated this pulldown in the CRISPR-edited cr_Q860X and observed similarly reduced interaction with the introduction of the C-terminal truncation (**Figure 4D**). Interestingly, we observed a higher level of ACTN1 in the unbound fraction of Q860X, both in the patient-derived hiPSC-CMs and cr_Q860X. Next, we analyzed ACTN1 levels in the cardiac tissue of the patient, and observed greatly elevated levels in both the left and right ventricle (**Figure 4E**), which could be a compensatory response to the dysfunctional interaction with ACTN2. It could also be indicative of the observed pathophysiology, as elevated ACTN1 protein levels have been observed in myocardium from failing hearts^44^. Despite elevated levels of ACTN1 in the patient, no interaction with ACTN2 was observed. For GJA1, we were unable to confirm interaction through Co-IP followed by Western Blot.

To further investigate the ACTN1-ACTN2 and GJA1-ACTN2 interactions in cardiomyocytes, we used double-label immunofluorescence in hiPSC-CMs followed by confocal microscopy imaging. The results showed a significant colocalization of GJA1 and ACTN1 with ACTN2 (**Supplementary Figure 12**). Thus, we have demonstrated several lines of evidence to support an interaction between ACTN2 and ACTN1, and ACTN2 and GJA1 in cardiomyocytes. No difference in the colocalization between either proteins was observed between Ctrl and Q860X in this analysis.

## Discussion

We show how protein-truncating variants in *ACTN2* causes cardiomyopathy with prominent arrhythmic phenotypes in the heterozygous state, and causes a progressive, severe restrictive cardiomyopathy in the homozygous state. Through deep cellular phenotyping and functional assays, we have demonstrated how these truncating variants cause disease and verified the causal nature through gene editing. Furthermore, our results have defined a protein-protein interaction network for alpha actinin 2 in human iPSC-derived cardiomyocytes, suggesting the importance of novel signaling pathways in the pathogenesis of *ACTN2*-mutation associated cardiomyopathies.

Alpha-actinin 2 is a highly conserved integral protein within skeletal muscle and cardiac sarcomeres^5^. The high-resolution three-dimensional structure of ACTN2 was recently revealed^6^. In *Drosophila*, lack of *ACTN2* is lethal^45^, and it is required for locomotion and proper assembly of Z-disc like dense bodies in *C. elegans*^46^. In a Zebrafish model, knockdown of *ACTN2* causes severe cardiac, skeletal muscle and ocular defects. Skeletal muscle manifestations included disorganized fibers, muscle weakness and overall immobility. Cardiac expression of *ACTN2* was observed already in the first stage of cardiogenesis, in both atria and ventricles, and mutant fish developed enlarged, dilated hearts. Importantly, this pathogenic phenotype could not be rescued by *ACTN3*, despite high structural similarity between the two isoforms^47^. In the patients described here, the lack of specific regions of ACTN2 similarly lead to cardiac disease and milder skeletal muscle manifestations.

The study of protein functional domains typically requires protein engineering. Here, we have utilized natural experiments to study the function of two domains in ACTN2. In the heterozygous state, we demonstrated the successful translation of transcripts from the allele carrying the large indel, and their incorporation into cardiomyocyte sarcomeres. The extent of the structural changes observed, with larger Z-discs, likely lead to altered contractile properties, where elevated Ca^2+^ signaling is a way for the cell to compensate for the contractile dysfunction. This establishes that protein-truncating *ACTN2* variants can have dominant negative effects causative of cardiomyopathy and arrhythmia.

The majority of genetic variants in sarcomeric genes that have been demonstrated to cause cardiac disease have been heterozygous. Our data show how a biallelic variant in *ACTN2* causes disease through a recessive model. *ACTN2* has a pLi score of 1; a complete intolerance to loss of function variation. However, this study describes a woman homozygous for a loss-of-function mutation in *ACTN2*, which presents an extremely rare case of a human C-terminal knockout. This patient has allowed us to study not only the pathogenic effect of this mutation, but also the physiological function of *ACTN2* and how lack of the C-terminal region disrupts the physiological role of ACTN2 in cardiomyocytes. The patient developed early-onset RCM with progression to end stage heart failure in her early 20s. Genes that have been associated with a restrictive phenotype are also associated with HCM and DCM, and many described families have members with each cardiomyopathy subtype. Examples include *MYBPC3^48^, TNNI3, TNNT2, ACTC* and *MYH7*^49–51^. Only one previous study has associated a likely pathogenic variant (N175Y) in *ACTN2* with RCM^52^. Our study is the first to show evidence for genetic variation in *ACTN2* as a cause of RCM through a recessive mode of inheritance.

Understanding underlying genetic causes of cardiomyopathy has major implications for diagnosis and screening of unaffected family members, and establishing the mechanisms that underpin disease pathogenesis is critical for development of targeted therapies. Multiple genetic variants in sarcomeric genes are well-established to cause different forms of cardiomyopathy, *e.g. MYH7, MYBPC3, TTN* and *LMNA.* There are several recent papers suggesting that only a core set of genes, not including *ACTN2*, are causative of cardiomyopathy^53, 54^. However, there are a growing number of publications that have associated genetic variants in *ACTN2* with cardiomyopathy^15–19^, as well as a regulatory element of *ACTN2* that was recently associated with heart failure^22^. We believe that the previous associations, in combination with the evidence presented here for how two protein-truncating variants, one dominant negative and one recessive, cause cardiac disease with variable pathological presentation supports inclusion of *ACTN2* as a disease-causing gene for cardiomyopathy in humans. Moreover, our successful modeling of *ACTN2* genetic disease using hiPSC-CMs presents an opportunity to identify more targeted therapies and test the potential alleviating effect of different drugs directly on myopathic cells.

## Materials and Methods

### Patient identification and genetic variant detection

Both probands underwent clinical panel genetic sequencing for genes associated with cardiomyopathy (GeneDx). Clinical laboratory adjudication for both variants was ‘variant of uncertain significance’. Site-specific testing for family members of patient 2 was performed by GeneDx and reported as ‘variant of uncertain significance’ based on ACMG guidelines. The heterozygous deletion was verified in patient-derived cells using standard Sanger sequencing after PCR amplification of the target region *(ACTN2* primers: CATCTCAGGGGAAAGGCTGC, TGCCTGAACTTCTCAGCCAG), where a 41bp insertion was also identified.

### Cardiac tissue sectioning and staining

Cardiac tissue sections from the explanted tissue were subjected to Hematoxylin and Eosin staining for cell size and Trichrome staining for detection of fibrosis. This was performed by the Stanford Medicine Human Histology Center, using standardized protocols.

### hiPSC culture and cardiomyocyte differentiation

Human iPSC lines for the healthy controls, Q860X and Del 1 were obtained from the Stanford University Biobank. The hiPSCs for Del 2 were obtained from the Coriell Institute (CW30314). All hiPSCs were maintained in Essential 8 medium and passaged until P20 before initiation of differentiation. Pluripotency of all hiPSC cell lines was confirmed with immunofluorescence staining for four pluripotency markers (Nanog, Oct4, SSEA3 and TRA-1-81, **Supplementary Figure 4**), see immunofluorescence methods for details. Differentiation into hiPSC-CMs was performed on a monolayer of cells using a small molecule Wnt-activation and inhibition protocol as previously described ^55^. All experiments were performed between day 30 and 45 post initiation of differentiation.

### Gene expression profiling and analysis pipeline

RNA was extracted from pulverized left or right ventricle human cardiac tissue using the *mix*Vana miRNA isolation kit (Thermo Fisher Scientific, #AM1560), according to the manufacturer’s specifications. RNA was isolated from cultured cells using the TRIzol method (Thermo Fisher Scientific, #15596026). RNA sequencing was performed on human cardiac left ventricle RNA from Q860X and three healthy control hearts and on hiPSC-CMs from all disease lines and two different healthy control lines. All sequencing was performed as paired end 2×150bp and generated an average of 30.9 million and 25.4 million uniquely mapped reads per sample for the cardiac tissue samples and cells respectively. In brief, adapter trimming was performed using Cutadapt and reads were aligned to the human reference genome (hg38) from Gencode (release 30, available at www.gencodegenes.org/human/) using STAR and counts were generated using the RSEM program. Patient FPKM values for each gene were compared to the corresponding healthy controls for each experiment (cardiac tissue or hiPSC-CMs). After removal of unexpressed genes that had FPKM < 1 in either group, fold differences (FDs), Z-scores, and differential expression significance of the 14,319 remaining genes were calculated using limma^56^. Gene set enrichment analysis was performed on the ranked fold differences using the fGSEA R package^57^. For qRT-PCR, 1-2 ug total RNA was reverse transcribed to cDNA using the High-Capacity cDNA Reverse Transcription kit (Thermo Fisher Scientific, #4368814). Real-time PCR was performed using specific primers for ACTN2 (GCTGAAGAAATTGTTGATGG, ATATCCTGAATAGCAAAGCG), with RPLP0 (AAGGCTGTGGTGCTGATG, CGGATATGAGGCAGCAGTTT) as a housekeeping gene, on a ViiA 7 Real-Time PCR System (Thermo Fisher Scientific) and quantified using the ΔΔC^T^ method.

### Protein isolation and immunoblot analysis

Total protein was isolated from whole or pulverized human left ventricle, right ventricle and skeletal muscle tissue. Each sample was mixed with RIPA buffer with 1x PI cocktail (Thermo Fisher Scientific, #78442). Cardiac tissue was homogenized with a bullet blender (4×1min, 3 minutes on ice between runs), while skeletal muscle was homogenized using a Dounce tissue grinder, while on ice. After centrifugation at 1500*g* for 10 minutes, the supernatant was used to assess protein concentration using the Pierce BCA Protein Assay kit (Thermo Fisher Scientific, #23225) before subsequent analysis. Cultured cells were washed once in PBS, incubated with ice-cold RIPA buffer + 1X PI cocktail for 10 minutes on ice, and snap-frozen in liquid nitrogen before protein quantification and analysis. For membrane protein isolation from hiPSC-CMs, cells were resuspended in TE buffer with 1X PI cocktail and centrifuged at 40,000 x *g*. After a repeated centrifugation, membranes were solubilized using Triton X-100 and subsequently centrifuged at 100,000 x *g* for 30 min before subsequent quantification and analysis.

Protein homogenates (10-20 ug) were separated on gradient TGX gels (Bio-Rad, #4568084 or #4568095) and blotted onto PVDF membranes. After blocking with Odyssey TBS blocking buffer (Li-Cor, #927-50000), the membranes were incubated with primary antibody (ACTN2: ab68167; ACTN1: ab68194, Abcam; GAPDH: sc-48167, Santa Cruz Biotechnology), and after washing, the corresponding IRDye fluorescent secondary antibody (Li-Cor). GAPDH was used as a loading control antibody on the same membrane. Immuno-complexes were imaged and quantified using the Li-Cor Odyssey Fc imaging system (Li-Cor).

### Immunofluorescence for cell size and pluripotency

Cells were washed once in PBS. hiPSC-CMs were then treated with 50mM KCl in PBS to relax the beating cells. After three subsequent washes in PBS, the cells were fixed in 4% Paraformaldehyde. After two washes in PBS, the cells were permeabilized with 0.1% Triton X-100, washed 3×5 minutes in PBS, and subsequently blocked with 5% goat serum for 30 min. Incubation with primary antibodies (1:25-1:1000, diluted in 5% goat serum) was performed overnight at 4°C, followed by 1x wash in PBS, 3×5 minutes gentle shaking washes in 0.1% Tween 20 in PBS, and a last wash in PBS. Cells were incubated with the appropriate secondary fluorescent antibody (diluted 1:1000 in 5% goat serum in PBS) for 1h at RT in the dark, followed by 1x wash in PBS, 3×5 minutes gentle shaking washes in 0.1 % Tween 20 in PBS, and a last wash in PBS (all dark). Cells were either coated in PBS or co-stained using probes for Actin (Invitrogen, #R37110) and/or Nucblue (Invitrogen, #R37606), and then coated in PBS until imaging. Cells fixed on glass were mounted onto glass slides using the Prolong Diamond Antifade Dapi5 (Thermo Fisher Scientific, #P369-62).

For analysis of cell size, cells were stained with a primary antibody against ACTN2 (Millipore Sigma, #A7811), and co-stained with probes for Actin and Dapi. Images were obtained at 10x magnification on a Leica DMI3000B (Leica Microsystems), and analyzed using Image J. Only cells positive for ACTN2, as a cardiomyocyte marker, were analyzed. All imaging and analysis were performed blinded.

Pluripotency of all hiPSC lines was verified through immunofluorescent staining for the pluripotency markers Nanog (Stemgent, #09-0020), Oct4 (Thermo Fisher Scientific, #MA1-104), SSEA-3 (Thermo Fisher Scientific, #MA1-020) and TRA-1-81 (Thermo Fisher Scientific, #41-1100) (**Supplementary Figure 4**).

### Contractility and Ca^2+^-handling analysis

For contractility analysis, hiPSC-CMs cultured on glass-bottomed dishes (MatTek) were placed on a heated platform (Warner Instruments) mounted on the stage of an inverted Olympus IX70 microscope. Two or three of spontaneously beating hiPSC-CMs were randomly selected in each dish, and the contractility analysis was performed in RPMI 1640 without glucose medium (Corning, #10-043-CV), supplemented with B27 (Thermo Fisher Scientific, #17504001) and lactate (Thermo Fisher

Scientific, #L7900) at 37°C. Contraction of the cells was measured at 240 Hz for 1 min using the edge-detection system (IonOptix) by tracing the displacement of the cellular edge. For each data, five or more twitches were averaged and contractility parameters were determined using IonWizard software (IonOptix). The contraction amplitude was expressed as a percentage change in cellular length.

For Ca^2+^-handling analysis, a similar experimental setup as for contractility was used. At approximately day 25 of differentiation, single iPSC-CMs from Q860X, Del1, Del2 and healthy controls were seeded at a low density on Matrigel (Corning, 1:10 dilution in PBS) coated rectangular 10 kPa micropatterns, with a 7:1 length:width aspect ratio as described by Ribeiro et al. ^58^, in glass-bottom dishes (MatTek). Between day 30-35, CMs were loaded with 2μM Fura2-AM (Thermo Fisher Scientific, #F1221) with 0.02% Pluronic F-127 (Thermo Fisher Scientific, #P3000MP) in 0 Ca^2+^ Tyrode solution for 8 minutes at room temperature as described previously^59^. CMs were washed twice and final [Ca^2+^] was brought up to 1.8 mM. Cells remained at room temperature for approximately 30 minutes to allow for dye de-esterification and then were mounted on the IonOptix system and field-stimulated at 1 Hz at 37°C. Single-cell calcium transients were measured at excitation wavelengths of 340 and 380 nm. Analysis and fitting of transient parameters was done using IonWizard software (IonOptix). Statistical analysis was performed using GraphPad (Prism).

### CRISPR-Cas9 gene editing to recapitulate Q860X

RNA guides were designed using CHOPCHOP (https://chopchop.cbu.uib.no/) and two guides cutting out the C-terminal region of *ACTN2* were selected (TCGGGAGCTGCCCCCGGATC and ATCGCTCTCCCCGTAGAGTG). Both primers were produced with annealing ends for the pX330 plasmid containing Cas9 (**Supplementary Figure 7**). After alignment of the sense and antisense oligos for each guide, the restriction enzyme *BbsI* was used to incorporate the guides into the pX330 plasmid. Following Sanger sequence verification, positive colonies were amplified and plasmids purified using a Qiagen Midiprep kit (Qiagen) according to the manufacturer’s specifications.

Human control iPSCs were cultured to 80% confluence. Cells were detached with Accutase (Sigma, #A6964) for 5 mins at 37°C. After washing in cell medium, cells were centrifuged at 300 RCF at RT and the cell pellet was subsequently resuspended in Buffer R from the Neon® transfection system kit (Thermo Fisher Scientific, #MPK10025) to a concentration of 40 million cells per ml. Nucleofection was performed on a Neon® Transfection System (Thermo Fisher Scientific, #MPK5000) on 5ul cells mixed with 1.5ug of each plasmid in 5ul Buffer R. The cell-DNA mix was exposed to one single pulse of 1100mV for 30s and subsequently replated on matrigel-coated plates (Corning) with Essential 8™ media (Thermo Fisher Scientific, #A1517001). After 2 weeks, multiple colonies were picked, replated and expanded. DNA was isolated using the QuickExtract kit (Epicentre, #QE09050) and the target region in the last exon of *ACTN2* (Primers: ATGTTGTGGTGTTTCTGCAACT, TATTGCATTCTGATGGGATGAG) was amplified and successful gene editing was verified through an agarose gel (**Supplementary Figure 7B**) and Sanger sequencing of the PCR product. Successfully edited cells were subsequently used for hiPSC-CM differentiation followed by phenotyping.

### NRVM isolation and analysis

Neonatal rat ventricular myocytes (NRVMs) were isolated from newly born rats using the neonatal rat/mouse cardiomyocyte isolation protocol from Cellutron (nc-6031) according to the specifications from the manufacturer. In brief, neonatal hearts were harvested from anesthetized newborn rats. NRVMs were extracted using multiple rounds of digestion with stirring at 37°C. On day 0, NRVMs were plated at a density of 0.2 million cells/well for 6 well plates, 0.5 million cells/well for 6 cm dishes, 2 million cells/plate for 10 cm plate in NS medium with 10% FBS plus glutamine. On day 1, the medium was changed to serum-free NW media and siRNA treatment was performed overnight using the Lipofectamine RNAiMAX Transfection Reagent (Invitrogen, 13778150) protocol. In brief, 30 pmol ACTN2 siRNA (Thermo Fisher Scientific, s145555) and 9ul Lipofectamine® reagent per well of a 6 well plate was each mixed with 150ul Optim-MEM I reduced serum media (Thermo Fisher Scientific, #31985062). After 5 minutes, the siRNA and lipofectamine solutions were mixed and incubated at RT for 20 mins before being added dropwise into the NRVM-containing wells. On day 2, the medium was changed to low-serum medium (0.2% FBS) for 48 hours for cell cycle synchronization. On day 4, the cells were harvested.

### Transmission electron microscopy

Cells were replated onto matrigel-coated 1cm x 5mm Aclar (Ted Pella, #10501-10). After 5 days of recovery, cells were relaxed using 50mM KCl and then fixed in Karnovsky’s fixative: 2% Glutaraldehyde (EMS, #16000) and 4% paraformaldehyde (EMS, #15700) in 0.1M Sodium Cacodylate (EMS, #12300) pH 7.4 for 30 min, chilled and sent to Stanford’s Cell Sciences Imaging Facility on ice. They were then post-fixed in cold 1% Osmium tetroxide (EMS, #19100) in water and allowed to warm for 2 hr in a hood, washed 3X with ultra-filtered water, then en bloc stained 2 hours in 1% Uranyl Acetate at RT. Samples were then dehydrated in a series of ethanol washes for 10 minutes each at room temperature beginning at 50%, 70%, 95%, changed to 100% 2X, then Propylene Oxide (PO) for 10 min. Samples are infiltrated with EMbed-812 resin (EMS, #14120) mixed 1:1, and 2:1 with PO for 2 hrs each. The samples were then placed into EMbed-812 for 2 hours opened then placed into flat molds w/labels and fresh resin and placed into 65°C oven overnight.

Sections were taken around 90nm, picked up on formvar/Carbon coated slot Cu grids, stained for 40 seconds in 3.5% Uranyl Acetate in 50% Acetone followed by staining in 0.2% Lead Citrate for 6 minutes. Observed in the JEOL JEM-1400 120kV and photos were taken using a Gatan Orius 2k X 2k digital camera. Quantification of sarcomere length, Z-disc thickness and I-band thickness was performed using ImageJ. For sarcomere length, an average of three measurements across each sarcomere was used, whereas for Z-disc and I-band thickness, the average of five measurements across each was used.

### ACTN2 heterozygous indel labelling

To visualize the ACTN2 protein product carrying the indel, we introduced the corresponding genetic sequence with a C-terminal FLAG-tag into a vector (**Supplementary Figure 9**). Differentiated control hiPSC-CMs were replated onto matrigel-coated (Corning) 35mm glass bottom dishes (MaTek). After recovery for 4 days, 5ug of the vector was introduced to the cells using the *Trans*IT-TKO transfection reagent (Mirus) according to the specifications by the manufacturer. A mock transfection without the vector was used as a negative control. The transfection was repeated after 24h to increase transfection efficiency. After an additional 24h, cells were fixed and stained with an anti-DDK antibody (OriGene, TA50011), and actin and dapi probes as described for immunofluorescence above. Confocal images at 40X magnification were obtained on an inverted Zeiss LSM 780 multiphoton laser scanning confocal microscope to visualize the FLAG-tagged protein within the hiPSC-CMs.

### Co-Immunoprecipitation followed by LC MS/MS

The Pierce™ classic magnetic IP/Co-IP kit (Thermo Fisher Scientific, #88804) was used for co-immunoprecipitation of ACTN2 in hipsc-derived cardiomyocytes from patient Q860X (3 replicates) and a healthy control sample (3 replicates). At day 36 post initiation of differentiation, the cell culture plates were placed on ice and the cells washed once in ice-cold PBS before addition of 100ul IP lysis/wash buffer (with 20ul/ml PI cocktail added fresh) was added to each well of a 12-well culture plate. After 10 min incubation on ice, the lysed cells were transferred to tubes and subjected to 10 min centrifugation at 13000xg. The supernatant was transferred into a fresh tube and protein concentration determined using the Pierce™ BCA protein assay kit (Thermo Scientific, #23225). 550ug protein from each sample was diluted to 500ul and mixed with 3 ug ACTN2 antibody (Abcam, ab68167) or 3ug normal rabbit IgG (Sigma-Aldrich, #12-370) as a negative control and incubated under slow mixing overnight at 4°C. The homogenate was subsequently incubated with magnetic beads for 1h at RT under constant mixing, and subsequently washed according to the manufacturer’s specifications. Samples were then re-suspended in ammonium bicarbonate and reduced with 10 mM DTT at 55°C for 5 min, followed by 25 minutes at room temperature. Alkylation was performed with 30 mM acrylamide for 30 min at room temperature. Digestion was performed with Trypsin/LysC (Promega) in the presence of 0.02% protease max (Promega) in a standard overnight digest at room temperature on head-over-head mixer. Digestion reaction was quenched with 1 % formic acid and peptides were de-salted with C18 Monospin reversed phase columns (GL Sciences). De-salted peptides were dried in a speed vac., reconstituted in reconstitution buffer (2% acetonitrile with 0.1% Formic acid) and 500ng of peptide material was injected on the instrument.

Mass spectrometry experiments were performed at the Stanford University Mass Spectrometry laboratory using an Orbitrap Q-Exactive HFX mass spectrometer (Thermo Fisher Scientific) with liquid chromatography performed using an Acquity M-Class UPLC (Waters Corporation). A pulled-and-packed fused silica C18 reverse phase column was used, with Dr. Maisch 1.8 micron C18 beads as the packing material and a length of ~25 cm. A flow rate of 450 nL/min was used with a mobile phase A of aqueous 0.2% formic acid and mobile phase B of 0.2% formic acid in acetonitrile. Peptides were directly injected onto the analytical column using a gradient (3-45% B, followed by a high-B wash) of 80min. The mass spectrometer was operated in a data-dependent fashion, with MS1 survey spectra and MS2 fragmentation collected in orbitrap using HCD.

Raw MS files were analyzed by MaxQuant (version 1.6.3.4)^60^, against a database containing 20,399 reviewed SwissProt human sequences^61^. All runs for a given bait were analyzed simultaneously to maximize the “match between runs” algorithm available on Maxquant. Multiplicity was set to 1 (as recommended for label-free experiments) and the false discovery rate imposed to 0.01 for peptide and protein identification. Statistical analysis of MaxQuant-analyzed data was performed with the artMS Bioconductor package^62^, which performs the relative quantification using the MSstats Bioconductor package^63^. Contaminants and decoy hits were removed. The samples were normalized across fractions by median-centering the log2-transformed MS1 intensity distributions. Log2(fold change) for protein/sites with missing values in one condition but found in >2 biological replicates of the other condition of any given comparison were estimated by imputing intensity values from the lowest observed MS1-intensity across samples^64^; p-values were randomly assigned between 0.05 and 0.01 to facilitate the inclusion of the imputed proteins when filters by p-value <0.05 were applied. SAINTq^65^ was used to assign scores to bait-prey interactions against the negative controls using peptide intensities as the input. Raw data, metadata, sequence file, and MaxQuant results files are available at the Mass Spectrometry Interactive Virtual Environment repository (MassIVE, https://massive.ucsd.edu/ProteoSAFe/static/massive.jsp) database.

Proteins illustrated in **Figure 4** were selected using the following criteria. For common identified proteins: FC>2 in both conditions compared to MOCK (IgG), p-value<0.02 in both, WT_CON_AvgP>0.95 (probability that it is a significant interactor in WT vs MOCK Control), Xpep1 (WT) ≥3 and Xpep2 (Q860X) ≥3 (number of peptides used for quantification in both WT and Q860X respectively). For proteins identified in WT but also significant in Q860X at less stringent p-value cutoff (<0.05): FC>2 in WT, p-value<0.02 in WT, p-value<0.05 in Q860X, WT_CON_AvgP>0.95, Xpep1 ≥3. For proteins identified in Q860X but also significant in WT at less stringent p-value cutoff (<0.05): FC>2 in Q860X, p-value<0.02 in Q860X, p-value<0.05 in WT, Q860X_CON_AvgP>0.95, Xpep2 ≥3. For proteins uniquely identified in WT: Significant difference between WT and Q860X (WT-Q860X_iLogFC>2 and WT-Q860X_iPvalue<0.05), FC>2 in WT, p-value<0.02 in WT, WT_CON_AvgP>0.95, Xpep1 ≥3, Q860X p-value>0.05 (not significantly identified in Q860X). For the identified proteins in WT, only LACTB remains, which is uniquely identified in WT, but not significantly different between WT and Q860X. For proteins uniquely identified in Q860X: Significant difference between WT and Q860X (WT-Q860X_iLogFC<-2, WT-Q860X_iPvalue<0.05), FC>2 in Q860X, p-value<0.02 in Q860X, Q860X_CON_AvgP>0.95, Xpep2 ≥3, WT p-value>0.05 (not significantly identified in WT). For the identified proteins in Q860X, only MYH11 and LRP2 remain, which are uniquely identified in Q860X, but not significantly different between WT and Q860X.

For the Western Blot validation analysis, the protein complexes were eluted from the magnetic beads in the elution buffer provided in the Pierce™ classic magnetic IP/Co-IP kit (Thermo Fisher Scientific, #88804) and subsequently analyzed according to the Western Blot protocol described above. Primary antibodies used were ACTN2 (Thermo Fisher Scientific, GT1253), ACTN1 (Abcam, ab18061) and GJA1 (Abcam, ab87645).

### Colocalization analysis

Beating hiPSC-CMs from Q860X and healthy control cells were replated onto matrigel micropatterns on 35mm glass-bottomed plates (MaTek) and allowed to recover for 4 days. The cells were subsequently fixed in 4% PFA and co-stained with ACTN2 (Thermo Fisher Scientific, A7811) and ACTN1 (Thermo Fisher Scientific, PA5-17308) or GJA1 (Abcam, ab87645), and a Dapi probe as described for immunofluorescence above. Confocal images at 40X magnification were obtained on an inverted Zeiss LSM 780 multiphoton laser scanning confocal microscope. Correlation Analysis: Confocal microscopy image stacks were analyzed using a custom image processing pipeline at the Stanford Shriram Cell Science Imaging Facility. The source code is available online at https://github.com/bioimage-analysis/malene_colocalization. This pipeline was created using algorithms from the scikit-image^66^, NumPy^67^ and SciPy^68^ Python packages. In order to create an ROI around the cell body, the signal from the green channel is first blurred using a gaussian filter with a sigma of 15. We then use the Li threshold algorithm to create a mask covering the cells which is further processed in order to remove small holes and small objects. The Pearson and Spearman correlation coefficients are then extracted from the ROIs.

## Acknowledgements

The authors would like to thank Dr. Alex Dainis for providing RNA from control hiPS cells from day 0 to day 10 of hiPSC-CM differentiation, and Kratika Singhal and Ryan Lieb at the Vincent Coates Foundation Mass Spectrometry Laboratory, Stanford University Mass Spectrometry for acquiring the LC-MS/MS data. This research was supported by the Knut and Alice Wallenberg Foundation (M.E.L) and in part by R01HL105993 from the National Institutes of Health (E.A.A). The project was supported, in part, by ARRA Award Number 1S10RR026780-01 from the National Center for Research Resources (NCRR). Its contents are solely the responsibility of the authors and do not necessarily represent the official views of the NCRR or the National Institutes of Health.

## Conflicts of interest statement

E.A.A. is founder at Personalis and DeepCell, Inc, and advisor for SequenceBio and Genome Medical. CC is an employee at GeneMatters. M.T.W. has ownership interest in Personalis. The other authors declare no competing interests.

