## Supplemental Figures for "Mono- and bi-allelic protein truncating variants in alpha-actinin 2 cause cardiomyopathy through distinct mechanisms"

### Supplementary figures

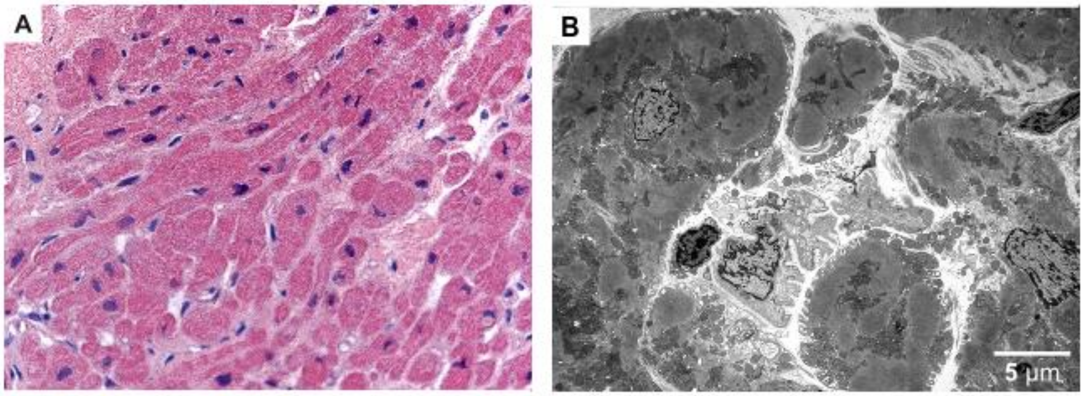


**Supplementary Figure 1.** Right ventricular endomyocardial biopsy obtained from patient Q860X at sixteen years of age demonstrating cardiomyopathic changes. Panel A: High power magnification showing myocyte hypertrophy characterized by enlarged, irregularly shaped myocyte nuclei. Fine interstitial fibrosis is noted between myocytes. (Hematoxylin & eosin x400); Panel B: Transmission electron microscopy of glutaraldehyde-fixed myocardium showing intact myocytes with preservation of mitochondrial distribution and alignment and contractile elements, variable nuclear alterations and absence of intracytoplasmic lipid accumulations. The microvasculature is normal (x2700).

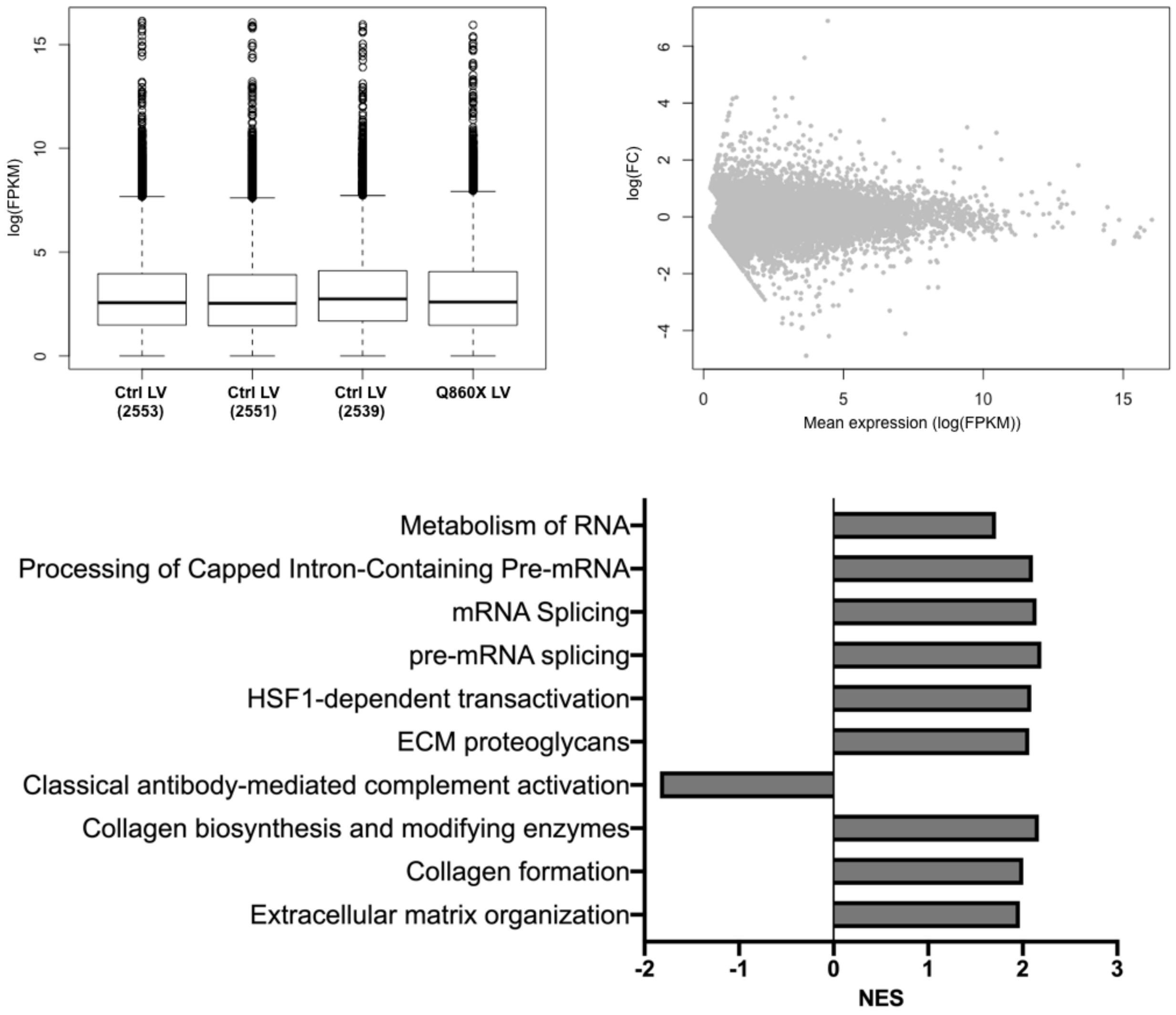


**Supplementary Figure 2.** RNA sequencing analysis of cardiac left ventricle tissue. Overall distribution of expression levels is similar between samples (left) and observed fold differences show the expected pattern (right). Gene set enrichment analysis (bottom). Significant reactome pathways (FDR<0.1) are displayed with Normalized enrichment scores (NES). N=3 controls.

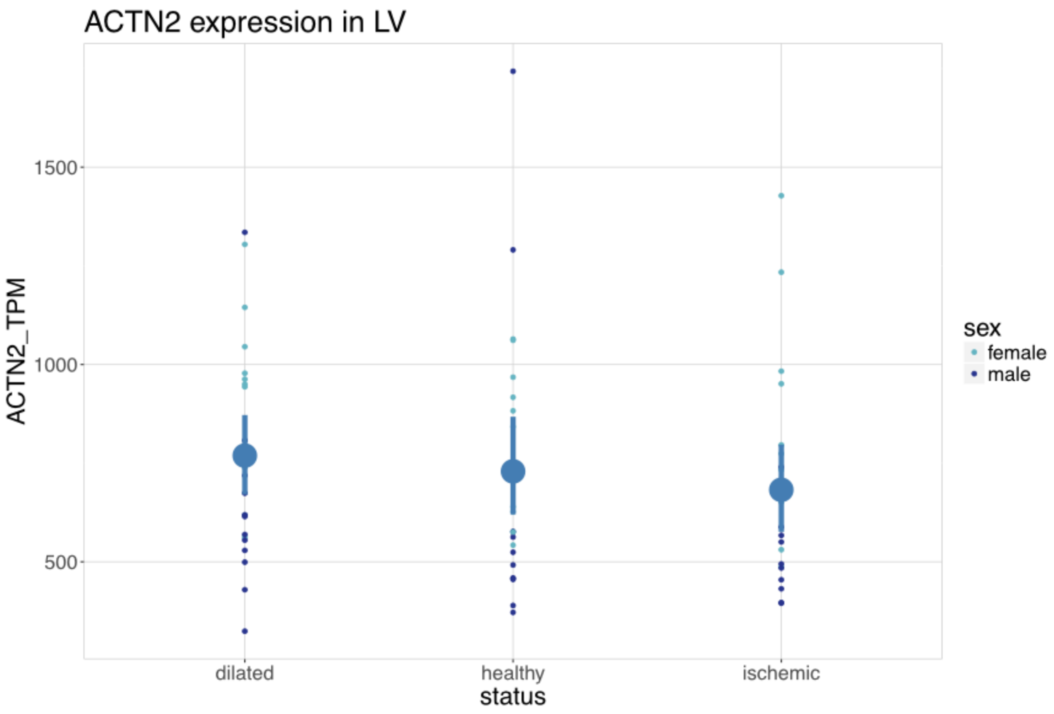


**Supplementary Figure 3.** Expression levels of ACTN2 in patients from the MAGNET consortium. Dilated cardiomyopathy (n=24), healthy controls (n=25) and patients with ischemic heart disease (n=21). Both male and female samples are represented.

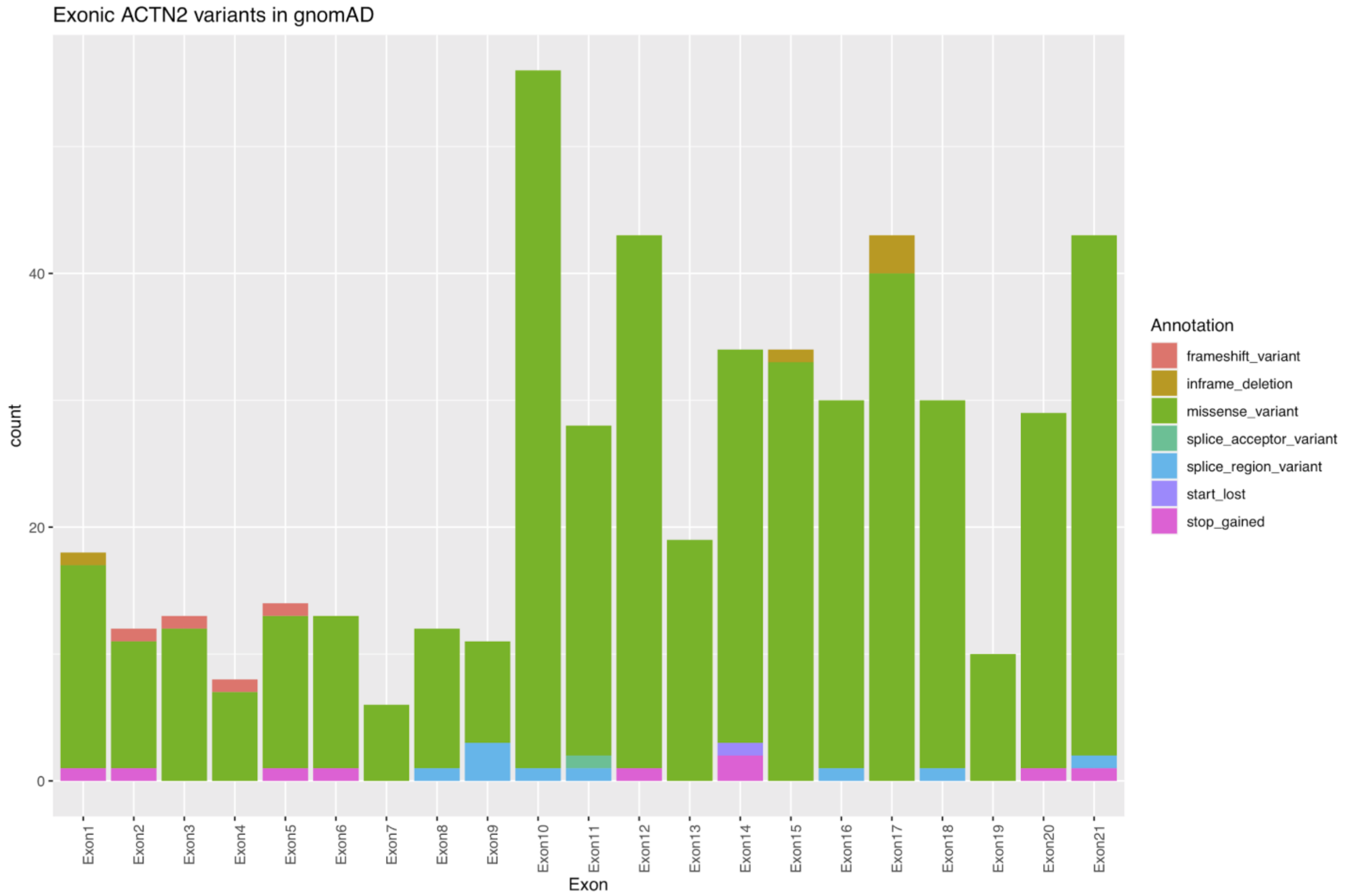


**Supplementary Figure 4.** Distribution of unique exonic variants in ACTN2 reported in GnomAD. Numbers are reported in actual counts.

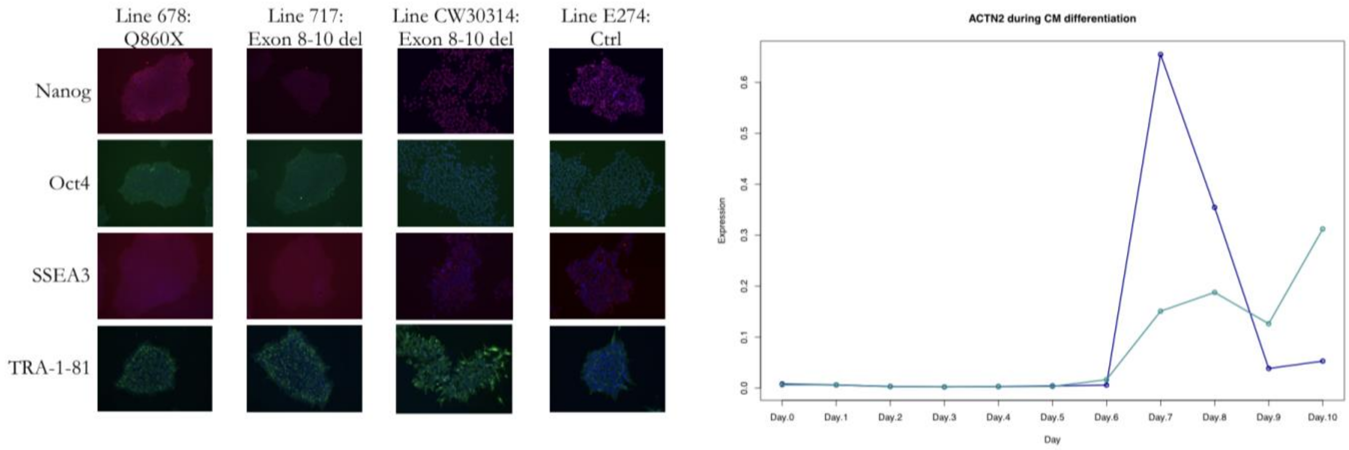


**Supplementary Figure 5.** Immunofluorescent staining of pluripotency markers Nanog in red, Oct4 in green, SSEA3 in red and TRA-1-81 in green, with nuclei stained with DAPI (blue) for the three patient hiPSC lines and one control line (left). Upon initiation of cardiomyocyte differentiation, *ACTN2* expression was observed around Day 6 post inititiation. Data shown for two different hiPSC lines.

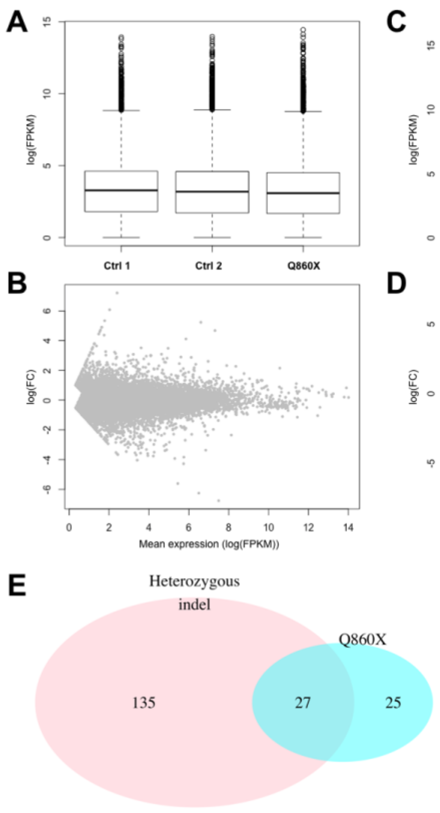


**Supplementary Figure 6.** RNA sequencing analysis of hiPSC-CMs. Overall distribution of expression levels between Ctrl samples (N=2) and Q860X (A) and observed fold differences in relation to mean expression level (B). Overall distribution of expression levels between Ctrl samples (N=2) and Del 1 and Del 2 (C) and observed fold differences in relation to mean expression level (D). E) Venn diagram of the pathways identified in the gene set enrichment analysis between the two *ACTN2* truncating variants.

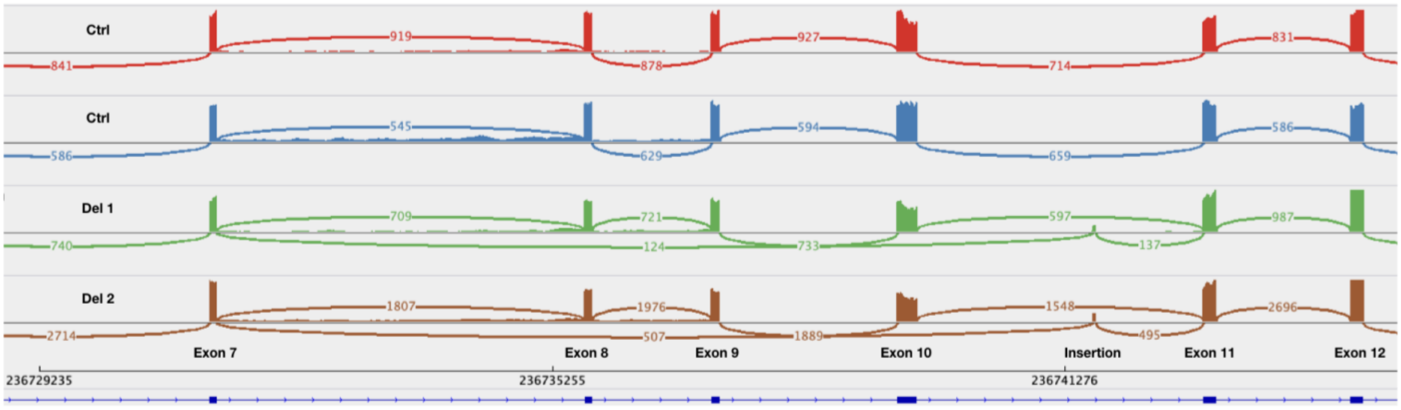


**Supplementary Figure 7.** Sashimi plot of the count data from RNA sequencing analysis of two control ipsc-CMs lines (red and blue lines) and patients Del 1 (green) and Del 2 (brown). Representation from Exon 7 to Exon 12 of the *ACTN2* gene.

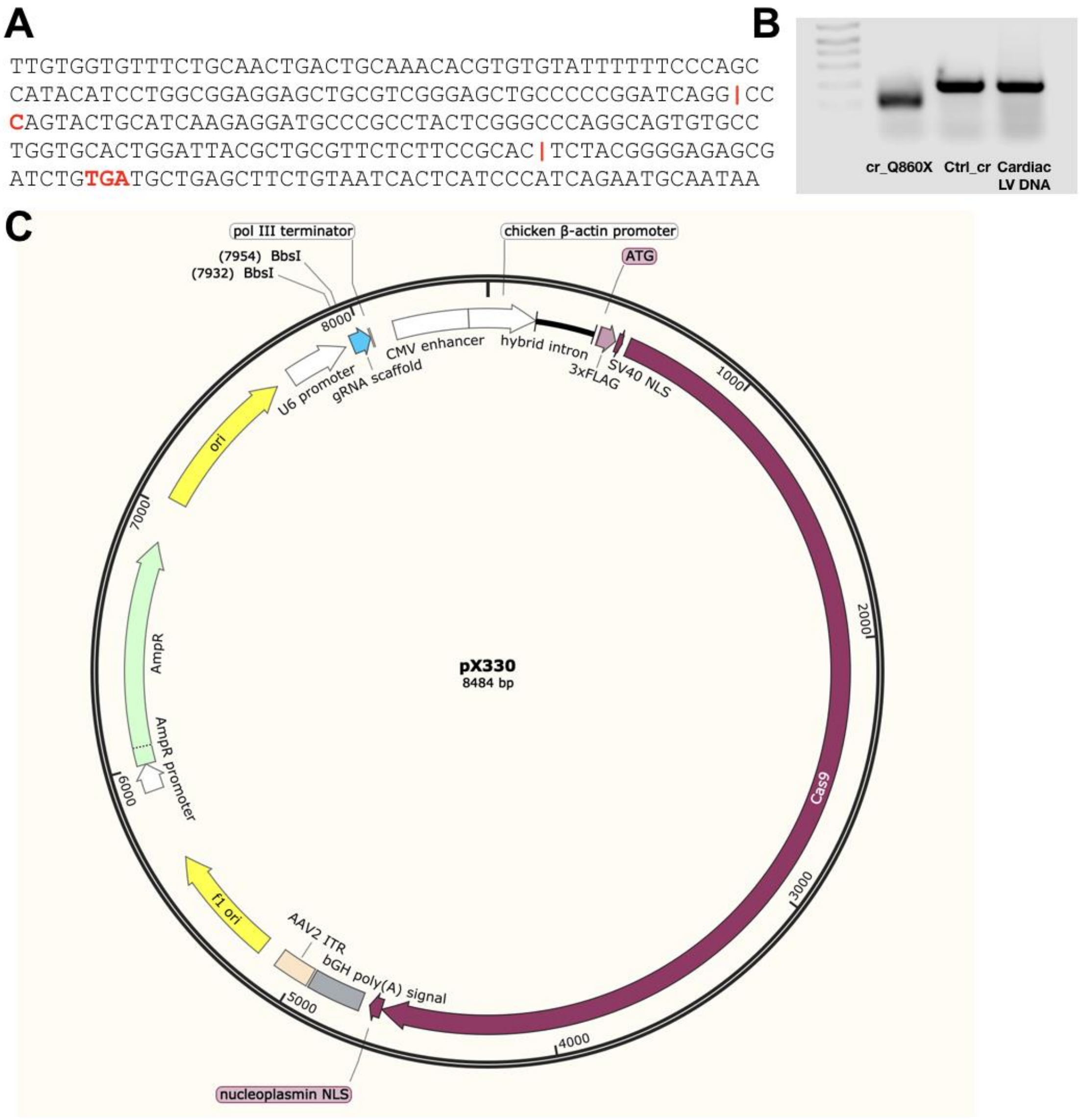


**Supplementary Figure 8.** CRISPR gene editing to simulate Q860X in healthy control hiPSCs. A) Guide RNAs were designed to cut out the same C-terminal sequence as in Q860X. The patient mutation site (c.2578C>T), the cut sites, and the new stop codon (TGA) are shown in red. B) PCR amplification of the region in the CRISPR:ed line (cr_Q860X), compared to control (Ctrl_cr) and a cardiac left ventricle control. C) The Cas9 plasmid used.

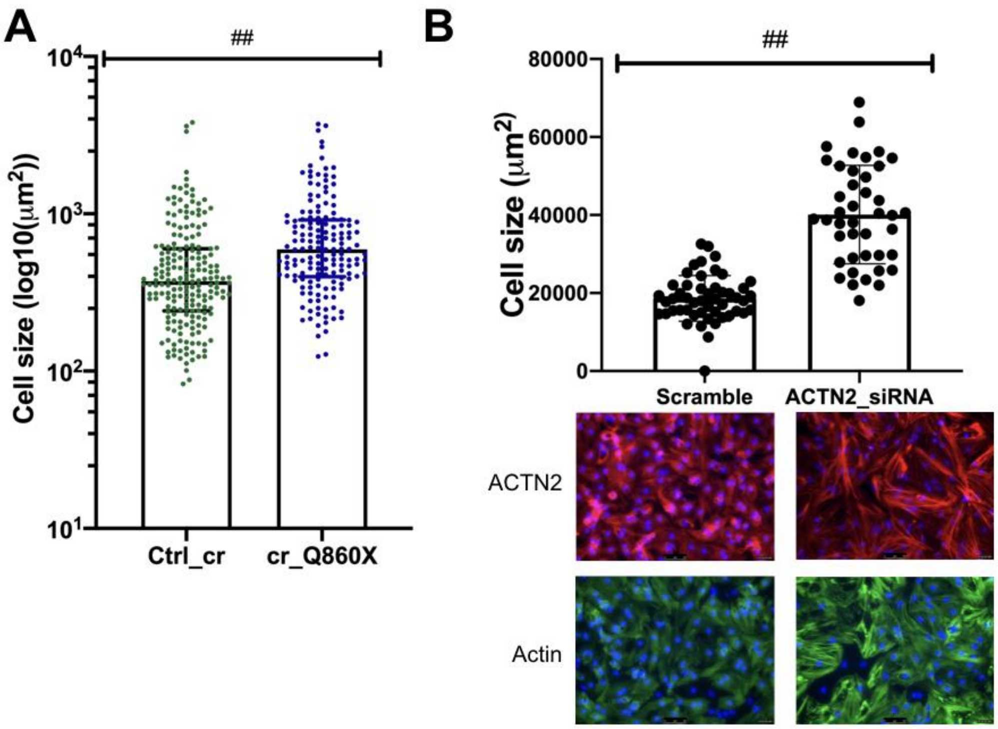


**Supplementary Figure 9.** Disruption of ACTN2 leads to cardiomyocyte hypertrophy. A) Cell size analysis comparing CRISPR-introduced C-terminal truncation of ACTN2 (cr_Q860X) with isogenic control hiPSC-CMs (Ctrl_cr). B) Cell size analysis of neonatal rat ventricular myocytes treated with siRNA to knock down ACTN2. Representative images of ACTN2 (red) and Actin (green) with Nuclei in blue (Dapi) are shown below each bar. ## indicates significant ($P$<0.001) based on a Kolmogorov-Smirnov test.

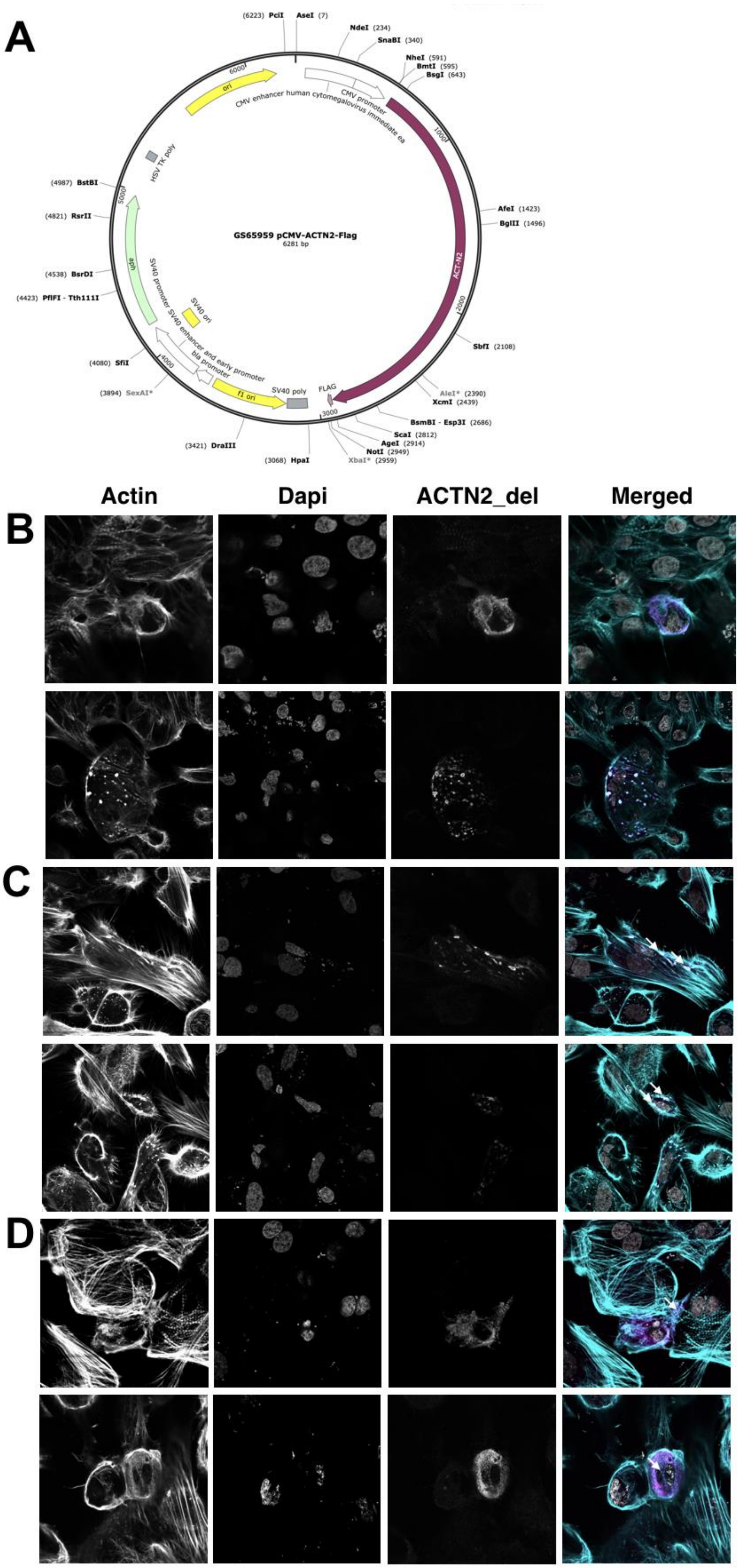
A
GS65959 pCMV-ACTN2-Flag
6281 bp
(6223) PciI
AseI (7)
NdeI (234)
SnaBI (340)
NheI (591)
BmtI (595)
BsgI (643)
AfeI (1423)
BglII (1496)
SbfI (2108)
AleI* (2390)
XcmI (2439)
BsmBI - Esp3I (2686)
ScaI (2812)
AgeI (2914)
NotI (2949)
XbaI* (2959)
(3068) HpaI
(3421) DraIII
(3894) SexAI*
(4080) SfiI
(4423) PflFI - Tth111I
(4538) BsrDI
(4821) RsrII
(4987) BstBI
ori
CMV enhancer human cytomegalovirus immediate ea
CMV promoter
ACT-N2
FLAG
SV40 poly
f1 ori
bla promoter
SV40 promoter SV40 enhancer and early promoter
SV40 ori
aph
HSV TK poly
Actin
Dapi
ACTN2_del
Merged
B
C
D

**Supplementary Figure 10.** Mechanisms heterozygous indel. A) Vector used to transfect FLAG-tagged ACTN2_del into hiPSC-CMs. B-D) Confocal images of transfected cells shown for Actin, Dapi, ACTN2_del and merged, where the merged is colored with Actin in turquoise, ACTN2_del in purple and Dapi in grey. B) shows how few cells were successfully transfected. C) Non-cardiomyocyte transfected cells typically displayed ACTN2 in droplets dispersed around the cell, as indicated by the white arrows. D) Successfully transfected hiPSC-CMs. Examples of sarcomeres displaying purple FLAG-stain are indicated by the white arrows.

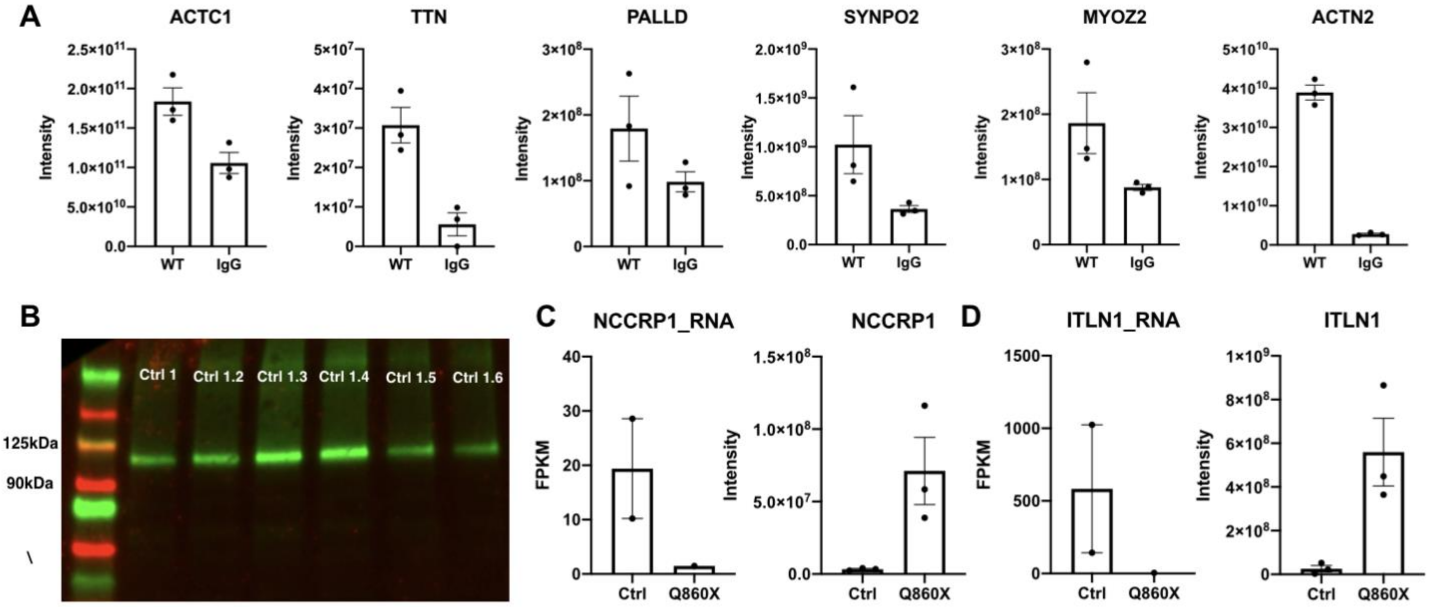


**Supplementary Figure 11.** The cardiomyocyte ACTN2 interactome. A) Intensity values from AP-MS data comparing WT ACTN2 pulldown with IgG negative control pulldown for ACTC1, TTN, PALLD, SYNPO2 and MYOZ2, that have all been previously shown to interact with ACTN2. ACTN2 itself is included for comparison. B) Ultra-centrifugation based membrane protein extractions of ipsc-CMs show abundant ACTN2 using Western Blot. C-D) RNA-seq (FPKM) and AP-MS data comparing control and Q860X for two genes from the interactome that showed differences in expression levels between control and Q860X; NCCRP1 (C) and ITLN1 (D).

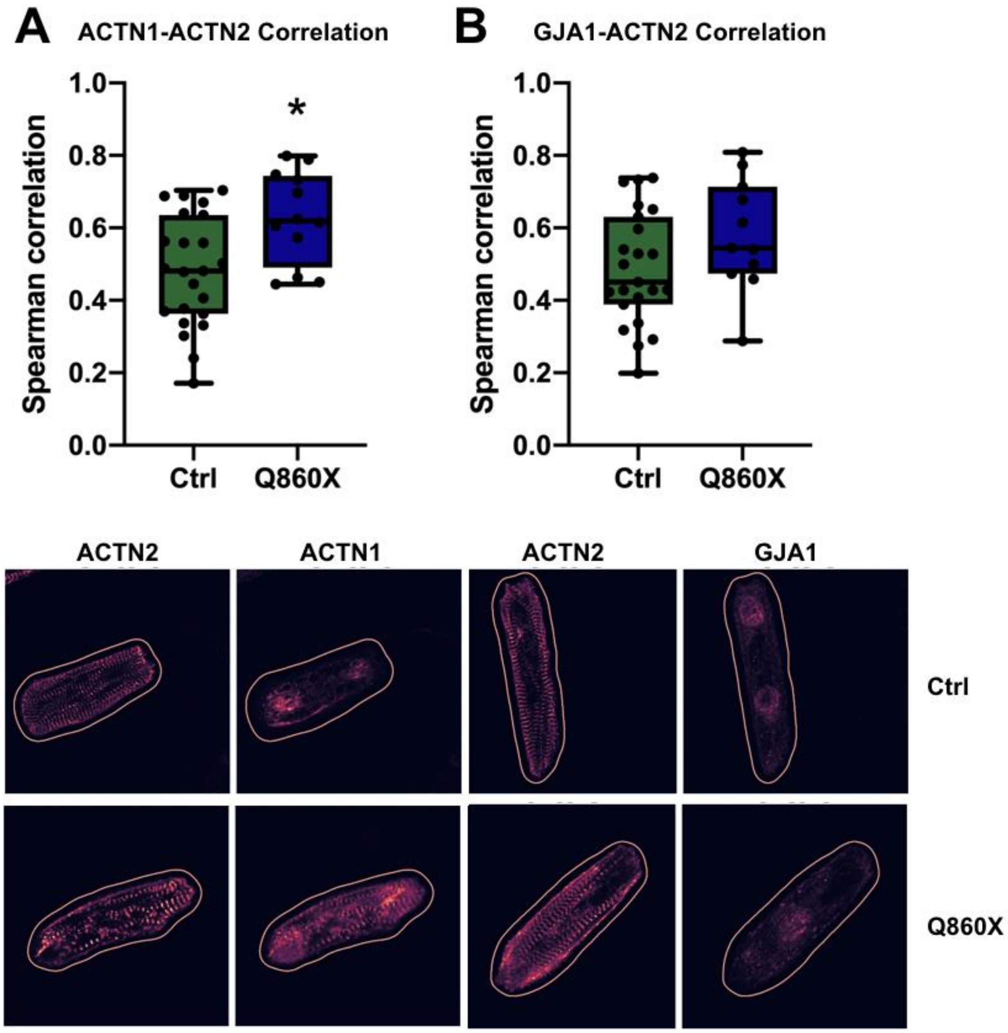


**Supplementary Figure 12.** Colocalization data. Confocal images of hiPSC-cardiomyocytes were analyzed for Spearman correlation between A) ACTN1 and ACTN2 and B) GJA1 and ACTN2. Representative images with regions of interest from the quantification are shown below each graph.
